# Unlocking new understanding of *Plasmodium* sporozoite biology with expansion microscopy

**DOI:** 10.1101/2025.04.09.648058

**Authors:** Benjamin Liffner, Thiago Luiz Alves e Silva, Elizabeth Glennon, Veronica Primavera, Elaine Hoffman, Alexis Kaushansky, Joel Vega-Rodriguez, Sabrina Absalon

## Abstract

Transmission of malaria relies on the formation of sporozoites in the mosquito midgut and their subsequent invasion of the salivary gland. Despite their importance, our understanding of the cell biology of sporozoite formation and salivary gland invasion is limited. Here, we apply a technique called <u>Mo</u>squito <u>Tiss</u>ue <u>U</u>ltrastructure <u>Ex</u>pansion <u>M</u>icroscopy (MoTissU-ExM), which physically expands infected mosquito tissues while preserving both host and parasite ultrastructure. Using MoTissU-ExM we are able to observe a range of parasite structures and organelles including features previously seen only by electron microscopy as well as structure not observed before. We leverage MoTissU-ExM to investigate a number of cell biology events during sporozoite formation and salivary gland invasion. In particular we focus on the rhoptries, a secretory organelle important for host cell invasion. We establish a timeline for sporozoite rhoptry biogenesis, show that two rhoptries are used up during salivary gland invasion, and provide the first evidence that rhoptry pairs are specialized for different invasion events. Building on this new knowledge, we characterize the rhoptry protein RON11 and identify it as the first protein involved in sporozoite rhoptry biogenesis. Disruption of RON11 led to the production of sporozoites that specifically fail to invade the salivary gland epithelial cell, thereby blocking transmission of these parasites.

## INTRODUCTION

Malaria parasites (*Plasmodium* spp.) have lifecycles that involve replication within and transmission between vertebrate and mosquito hosts^1^. Parasites transmitted to their mosquito host first undergo sexual reproduction^2^ before invading the mosquito midgut to form an asexual replicative lifecycle stage called an oocyst on the basal side of the midgut (Figure S1). In a single oocyst, thousands of sporozoites are formed in a process called sporogony^3,4^. Once mature, these sporozoites egress from the oocyst into the mosquito haemocoel, before entering the salivary glands (SG) of the mosquito (Figure S1). To do so, sporozoites cross the SG basal lamina, invade a SG epithelial cell, and move into the secretory cavity^5^; the mosquito anatomical structure that contains saliva (Figure S1). From the secretory cavity, sporozoites can be transmitted to a vertebrate host during a blood meal.

Little is known about the cell biology of sporozoite formation and function, particularly when it comes to essential cellular processes like mitosis, cytokinesis, or organelle biogenesis. The limited data that does exist largely comes from electron microscopy studies now decades old^6–8^, with relatively few functional reports on gene deleted or disrupted parasites. Oocysts initially undergo a growth phase over a few days, undergoing rounds of mitosis and rapidly increasing the number of genome copies contained in the oocyst^9^. The parasite plasma membrane then undergoes an invagination to form a structure called the sporoblast. Sporozoites will then form around the sporoblast, beginning to build their apical secretory organelles (rhoptries, micronemes, dense granules)^9^. Sporozoites are produced through a specialized cytokinesis process called segmentation^10,11^, defined by a contractile ring structure called the basal complex at the leading edge of the parasite. At the completion of segmentation, sporozoites bud off from the sporoblast and egress from the oocyst into the haemocoel.

From the haemocoel, sporozoites migrate to and invade the SG. Although some progress has been made, the molecular mechanisms underlying SG are poorly understood. Sporozoites cross the SG basal lamina and invade first the SG epithelial cell and subsequently secretory cavity; forming a vacuole after each invasion event. It has recently been shown that invasion of the salivary gland requires the secretion of proteins from secretory organelles called rhoptries^12–15^. Additionally, this invasion event has been shown to involve the formation of a tight-junction^12^, similarly to invasion of malaria parasites into red blood cells or hepatocytes. To date, only a handful or proteins have been implicated in SG invasion, highlighting a critical gap in our understanding of the molecular underpinnings of this essential step in malaria parasite transmission.

In the last few years, a technique called ultrastructure-expansion microscopy (U-ExM) has revolutionized the study of parasite cell biology^16^. U-ExM physically enlarges a sample ∼4.5-fold while preserving cellular ultrastructure^17,18^ therefore making ultrastructural analysis amenable with light microscopy. We have recently adapted U-ExM for use with dissected mosquito tissues, a technique we termed MoTissU-ExM^19^. Here, we apply MoTissU-ExM to investigate a range of cell biology processes during sporozoite formation within the oocyst, and then visualize the sporozoite’s invasion of the SG. Further, we investigate these processes across *P. yoelii*, *P. berghei*, and *P. falciparum*, the three *Plasmodium* sp. most commonly used to investigate mosquito-stage biology.

We show that using MoTissU-ExM, a range of oocyst structures and organelles can be visualized that have not been well studied previously. These include the apical polar ring (APR), basal complex, rhoptries, centriolar plaque (CP), rootlet fibre, and apicoplast. Due to the small size of the spindles that coordinate oocyst mitosis, this process has not been studied in detail. Using MoTissU-ExM to visualize both microtubules and the CP, which enables formation of intranuclear microtubules, we provide the first observation that the three microtubule spindle structures that coordinate mitosis in blood-stage malaria parasites, also do so in oocysts. We were also able to observed lateral asymmetry in sporozoites, which previously could only be seen using electron microscopy^20^. By imaging infected SGs, we were able to differentiate sporozoites at each of their stages of SG invasion, enabling us to examine perturbations in this process.

Much of this study focusses on the secretory organelles the rhoptries, which are secretory organelles that contain proteins that facilitate host cell invasion^21–23^. Despite their importance for SG invasion, little is known about the biogenesis in the oocyst or how they function during SG invasion. It is known that rhoptries golgi-derived, made *de novo* in oocysts, and that they are associated with a rhoptry-like structure called an AOR or rhoptry anlagen^6–8,24^. Beyond this, little is known about the process of rhoptry biogenesis, when it begins, and when it ends. We had recently used U-ExM to visualize rhoptry biogenesis in blood-stage malaria parasites^25^. In this study, we adopt a similar approach to establish a timeline for the biogenesis of rhoptries in sporozoites. Subsequently, we perform the first detailed quantification of rhoptry number during the sporozoite’s journey into the secretory cavity of the SG.

Having described a timeline for rhoptry biogenesis, we wanted to leverage this new understand to characterize the function of a rhoptry biogenesis protein. To do this, we investigated the protein RON11, which was known to localize to the rhoptries of sporozoites^14^ and be important for SG invasion^14^. Further, knockdown of RON11 in merozoites was recently shown to lead to the formation of merozoites with only a single rhoptry^26^. We show that stage-specific disruption of RON11 leads to a formation of sporozoites with rhoptry malformations that only contain half the number of rhoptries of controls. Additionally, we show that RON11 disrupted parasites are able to cross the SG basal lamina but fail to invade the SG epithelial cell, instead accumulating in the space between cells of the SG. Collectively, this study provides new insight into a range of cell biology processes during sporozoite formation in oocysts and SG invasion and highlights the utility of MoTissU-ExM for characterising perturbations in mosquito-stage malaria parasites.

## RESULTS

### Mosquito tissue Ultrastructure Expansion Microscopy (MoTissU-ExM) enables visualization of multiple parasite organelles and structures

A range of parasite structures and processes are visible using MoTissU-ExM^19^, but these have not previously been described in detail. Using MoTissU-ExM on *P. yoelii*-infected tissues in combination with fluorescent dyes for protein density (NHS ester Alexa Fluor 405), DNA (SYTOX deep red), or lipids (BODIPY-Fl-Ceramide), we were able to visualize a wide range of parasite structures and organelles without the need for protein-specific antibodies (Figures 1 and 2). Oocysts undergoing mitosis, contain large nuclei with multiple centriolar plaques (CPs) (Figure 1b) that presumably contain multiple genome copies, as previously reported^27^. Additionally, in mitotic oocyst nuclei, we visualized all of the microtubule spindle structures that are present in blood stage parasites^25,28–30^, the hemispindle, mitotic spindle, and interpolar spindle (Figures 1a and 1b). The early stages of sporozoite formation were observed through the formation of the apical polar rings (APRs); although the APRs themselves could not be readily distinguished from each other (Figure 1c). Once sporozoite formation has begun, the rootlet fiber that connects the CP to the apical end of the forming sporozoite^7,8,31^ is visible (Figure 1c). Once cytokinesis has begun the basal complex could be visualised, enabling determination of the progress of sporozoite formation through segmentation (Figure 1d). Rhoptries, along with the putatively rhoptry-associated organelle, known as an AOR^27^ or rhoptry anlagen^9,32,33^ (referred to as AOR hereafter), could be seen by protein density (Figure 1c). Occasionally, the sporozoite apicoplast could be observed by protein density alone, which we confirmed by staining with antibody markers of the apicoplast (Figure 1e).

**Figure 1:**
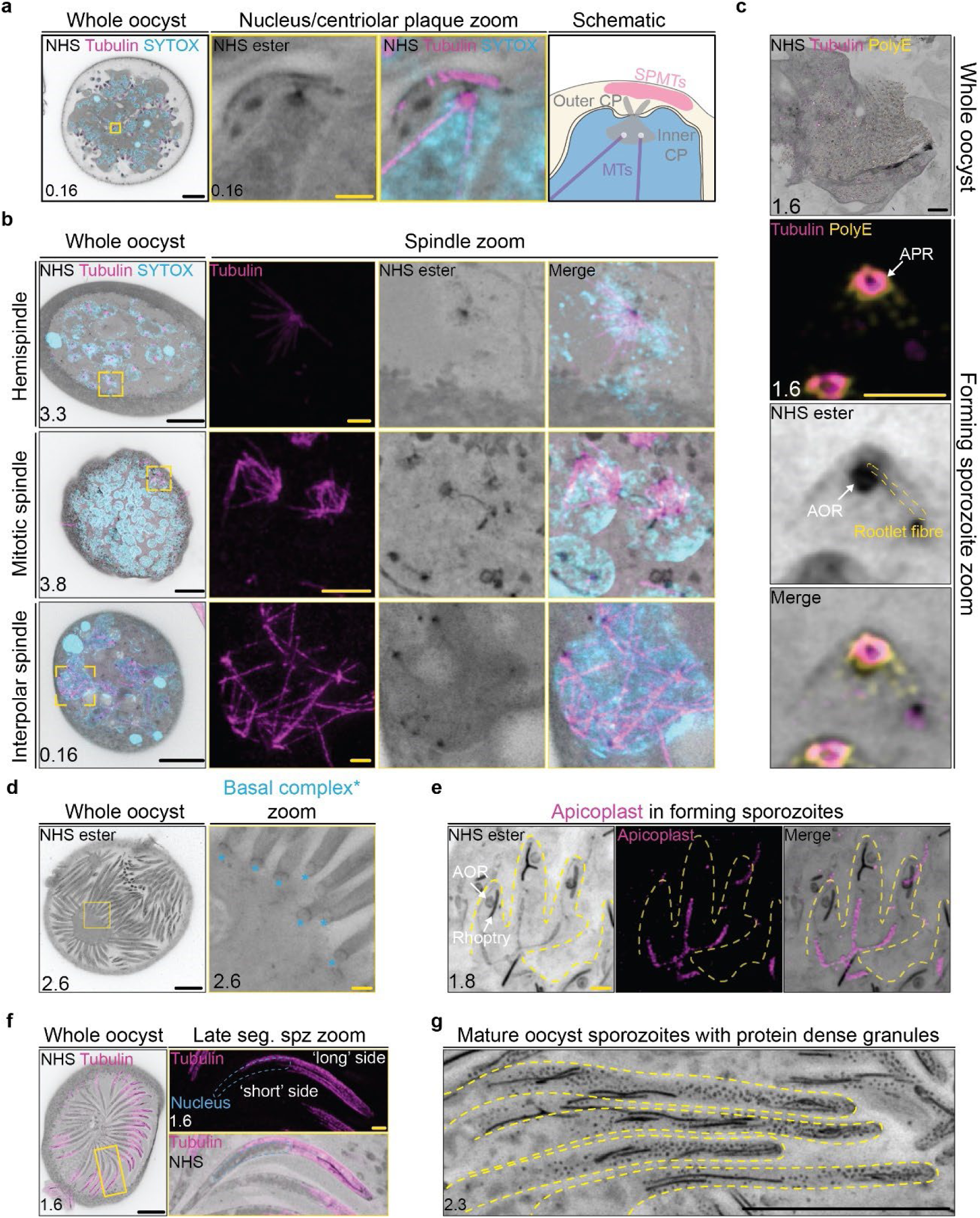
MoTissU-ExM enables visualization of a range of organelles and structures in oocysts and forming sporozoites. *Plasmodium yoelii* infected midguts were harvested on day 10 post infection, prepared by MoTissU-ExM, stained with the protein-density dye NHS Ester (greyscale), SYTOX Deep red (DNA, cyan) and other dyes, before being imaged by Airyscan2 microscopy. (**a**) Visualisation of the oocyst centriolar plaque (CP) and its associated microtubules (anti-tubulin, magenta). SPMTs = subpellicular microtubules. (**b**) CP and microtubules of oocyst nuclei at different stages of mitosis. (**c**) The apical polar ring(s), rootlet fibre, and CP of an early forming sporozoite stained with microtubule (anti-tubulin, magenta) and SPMT (anti-PolyE, yellow) markers. (**d**) The basal complexes of forming merozoites were visible by protein density, indicated by asterisks. (**e**) The partially segmented apicoplast (anti-acyl carrier protein, magenta) shared between multiple forming sporozoites. (**f**) Lateral asymmetry in a mature oocyst sporozoite visible through the extension of SPMTs (anti-tubulin, magenta) further towards the basal end of the sporozoite on the ‘long side’ compared to the ‘short side’. (**g**) Highly protein dense granules present in mature oocyst sporozoites. Scale bars: Black = 10 µm, Yellow = 2 µm. Number in bottom left corner indicates image Z-depth in µm.

**Figure 2:**
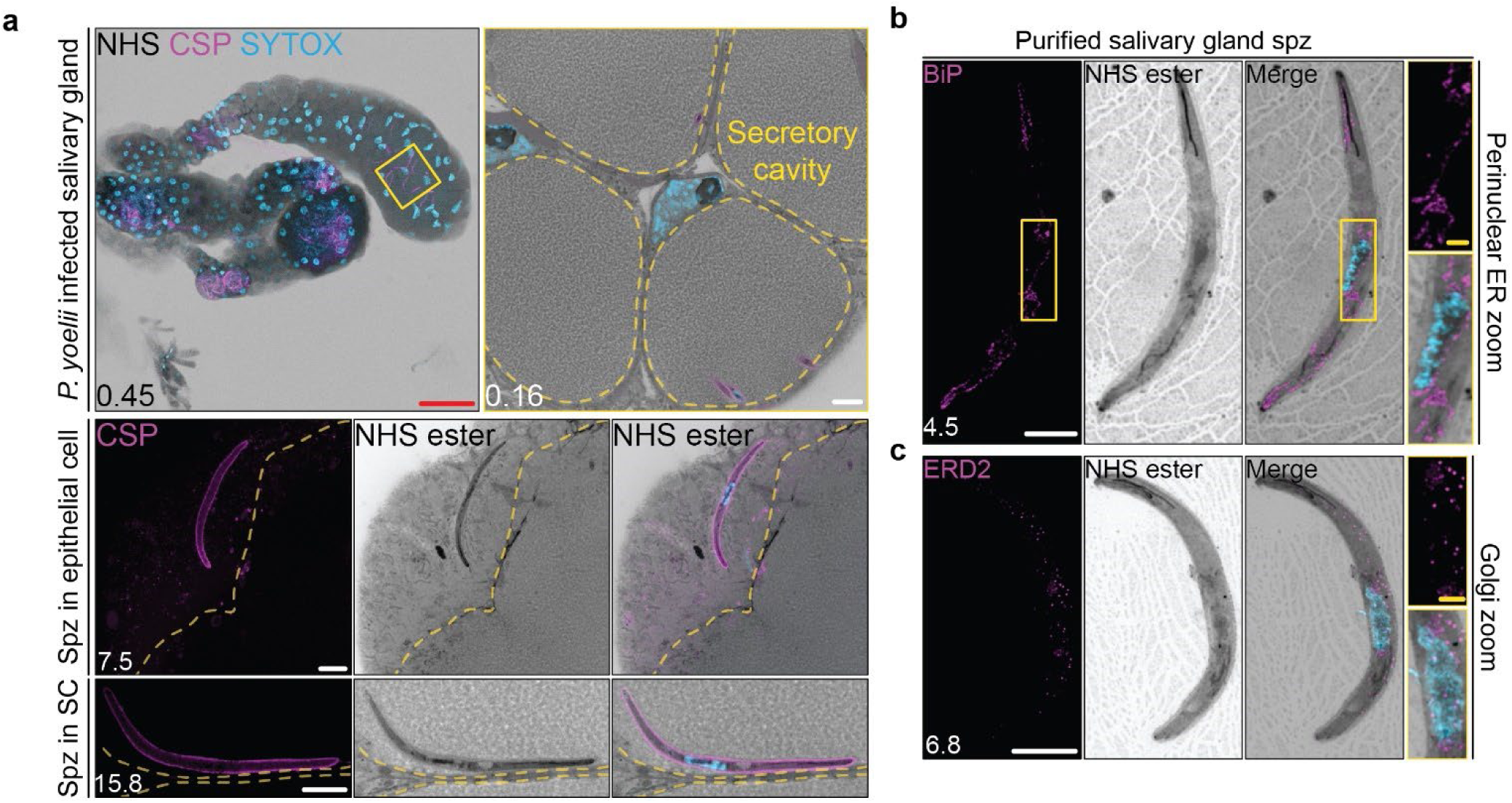
MoTissU-ExM enables visualization of sporozoites at different stages of salivary gland invasion. (**a**) *Plasmodium yoelii* infected SGs were harvested on day 14 post infection, prepared by MoTissU-ExM, stained with the protein-density dye NHS Ester (greyscale), SYTOX Deep Red (DNA, cyan) and anti-circumsporozoite protein (CSP, sporozoite surface, magenta), before being imaged by Airyscan2 microscopy. Sporozoites (spz) within SG epithelial cell and the secretory cavity (marked by yellow dashed line) could be differentiated. (**b**-**c**) Purified *P. yoelii* SG sporozoites stained with antibodies against either the endoplasmic reticulum marker BiP (**b**) or Golgi marker ERD2 (**c**). Scale bars: red = 200 µm, white = 10 µm, yellow = 2 µm. Number in bottom left corner = image z-depth in µm.

Sporozoite lateral asymmetry has previously been described using EM for the position of the nucleus, which is more closely associated with one side of the sporozoite plasma membrane than the other^20^. In mature oocyst sporozoites, or later in the lifecycle, we could observe this nuclear asymmetry (Figure 1f). Additionally, by staining expanded sporozoites with anti-tubulin antibodies, sporozoite subpellicular microtubules (SPMTs) could be seen displaying lateral asymmetry (Figure 1f). On the side of the sporozoite where the nucleus contacts the parasite plasma membrane (short side) the SPMTs were visibly shorter than the other side, where the SPMTs extended past the nucleus (long side). These observations show that sporozoites have laterally asymmetrical SPMTs and suggest that potentially this asymmetry is established early in sporozoite development.

Finally, we found that the cytoplasm of mature oocyst sporozoites contain a large number of highly protein-dense granules (Figure 1g). It is currently unclear what these granules are, or whether all these granules represent the same structure, but these granules seem to contain circumsporozoite protein (CSP) (Figure S2c) and their number reduces dramatically upon SG invasion (Figure S2d). Curiously, SG sporozoites typically contained two distinguishable layers of CSP, one directly at the sporozoite plasma membrane and a second, more distal, CSP layer encompassing the entire sporozoite (Figure S2b). The distal second CSP layer was present in haemolymph sporozoites (Figure S2c) and not associated with lipid staining using the lipid dye BODIPY Ceramide (Figure S2b), suggesting that it does not represent a vacuole isolated with the sporozoite. Whether there is any functional significance of this second CSP layer is currently unclear.

### Stage of salivary gland invasion by sporozoites can be visualized using MoTissU-ExM

Invasion of sporozoites into the SG is very poorly understood. After egressing from the oocyst, sporozoites are carried by the haemolymph flow to the SGs. To reach the SG secretory duct, where sporozoites will reside until transmitted to a vertebrate host, sporozoites must cross the basal lamina, invade a SG epithelial cell, and exit the cell while retaining cell viability^34^. Using MoTissU-ExM, we could observe fully mature oocyst sporozoites (Figure 1) and sporozoites either in the SG epithelial cell or secretory cavity (Figure 2a). This distinction between epithelial cell and secretory cavity enables a more detailed visualization of the progress of sporozoites through SG invasion. Further, we can use U-ExM to visualize isolated haemolymph, oocyst, or SG sporozoites. By imaging sporozoites purified from SGs in combination with organelle specific antibodies, we were able to identify both the sporozoite endoplasmic reticulum (Figure 2b) and Golgi (Figure 2c).

### Progress through cytokinesis as a proxy for oocyst developmental stage

*P. yoelii* oocysts were substantially different in their progress through sporozoite development, even within the same midgut (Figure 3a). In blood-stage *P. falciparum,* we have previously used progress through merozoite segmentation (cytokinesis) as a proxy for parasite development^25^. We adopted the same approach to develop a sporozoite segmentation score (Figure 3b). Using this metric, scores range from 1 (an oocyst that is still undergoing mitosis and has not yet begun segmentation) to 7 (an oocyst that contains morphologically mature sporozoites that have detached from the sporoblast), with each score in-between defined by a visible cell biology event as follows: 1 – oocyst still undergoing mitosis, 2 – apical polarity established and APR visible, 3 – apical heads of forming sporozoites visible, 4 – basal complex constricts the nucleus of forming sporozoite, 5 – basal complex past the nucleus of forming sporozoite, 6 – sporozoite appears morphologically mature but basal complex has not fully contracted, 7 – basal complex fully contracted, sporozoite detached from sporoblast (Figure 3b). Examples of oocysts at each of these segmentation scores across *P. yoelii*, *P. berghei*, and *P. falciparum* can be found in Figure S3. Using this metric for sporozoite development, we could approximately age-match oocysts to determine when particular events or processes occur either between different oocysts, or between different treatments.

**Figure 3:**
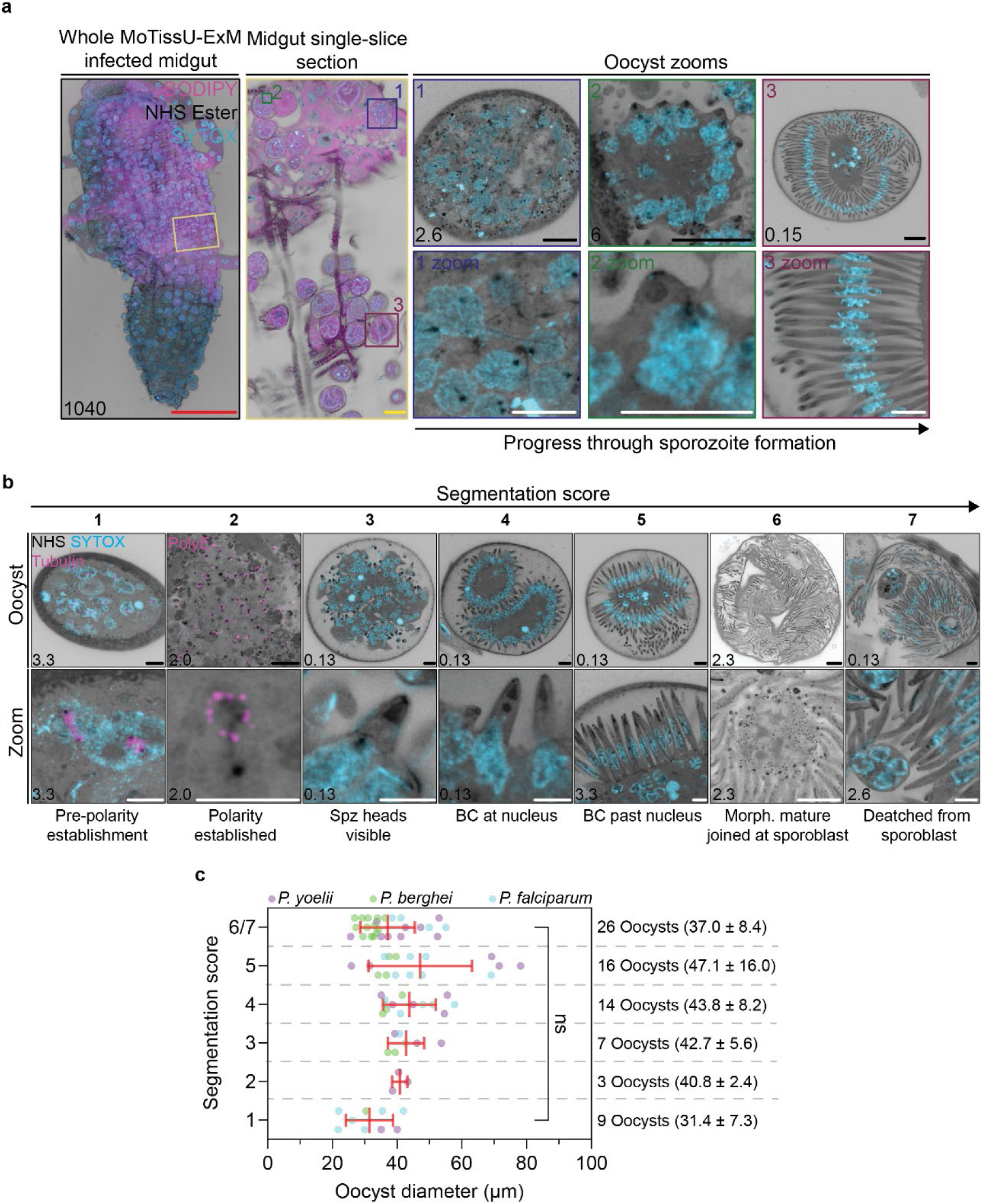
Using progress through segmentation as a readout for oocyst development. (**a**) *Plasmodium yoelii* infected midguts were harvested on day 15 post infection, prepared by MoTissU-ExM, stained with the protein-density dye NHS Ester (greyscale), SYTOX Deep Red (DNA, cyan) and lipid dye BODIPY FL ceramide (membranes, magenta), before being imaged by Airyscan2 microscopy. Images depict the whole infected midgut, a region of interest within that midgut (single-slice), and three oocysts within the region of interest at markedly different stages of sporozoite formation. (**b**) By imaging *P. yoelii* oocysts across the entirety of sporozoite formation a segmentation score, defined by the visualization of key biological events, was developed. Images are arranged in inferred order of sporozoite development. Spz = sporozoite. BC = basal complex. PolyE = anti-PolyE antibodies (subpellicular microtubules). Scale bars: red = 1 mm, yellow = 100 µm, black = 20 µm, white = 10 µm. Number in bottom left corner = image z-depth in µm. (**c**) In midguts infected with either *P. yoelii* (purple), *P. berghei* (green), or *P. falciparum* (cyan) the maximum diameter of oocysts were measured and their stage of segmentation determined (segmentation score). Segmentation scores of 6 and 7 were combined because separation of sporozoites form the sporoblast could not always be determined. Note that diameter measurements have been expansion factor corrected. 75 total oocysts were measured (*Py* = 27, *Pb* = 23, *Pf* = 25) across 15 midguts (*Py* = 5, *Pb* = 6, *Pf = 6*). ns = p>0.05 by ANOVA. Error bars = SD.

#### Oocyst diameter poorly reflects developmental stage during sporozoite formation

In the past, oocyst development has typically been inferred by measuring the size of oocysts^35^. Recent studies have shown that while oocysts increase in size initially, their size eventually plateaus (11 days post infection (dpi) in *P. falciparum*^4^). In oocysts where sporozoite formation had begun (segmentation score ≥2) we noticed little association between oocyst size and developmental stage. We measured the diameter of 75 oocysts from either *P. yoelii*, *P. berghei*, or *P. falciparum* and observed no statistically significant change in oocyst diameter between segmentation scores 1 and 6 (Figure 3c). Additionally, there was no correlation between oocyst diameter and segmentation score (r^2^=0.004; Figure S4). Our data therefore suggest that the plateau in oocyst size is coincident with the onset of cytokinesis. Collectively, this suggests that oocyst diameter is a poor readout for sporozoite development and highlights the usefulness of the segmentation score as an alternative.

### Defining a timeline for sporozoite rhoptry biogenesis

Rhoptries can readily be observed using the protein density stain NHS Ester, without the need for rhoptry-specific antibodies, as they are the most protein-dense structures in the parasite^25,36,37^. By staining expanded blood-stage *P. falciparum* with NHS Ester, we recently used progress through merozoite segmentation to stage parasites and define a timeline for merozoite rhoptry biogenesis^25^. Similarly using *P. yoelii* oocysts, we defined the first timeline for sporozoite rhoptry biogenesis (Figures 4a and 4b). We imaged oocysts across the entirety of sporozoite development, from before apical polarity establishment to sporoblast detachment, observing the rhoptries and how they change over time (Figure 4a).

**Figure 4:**
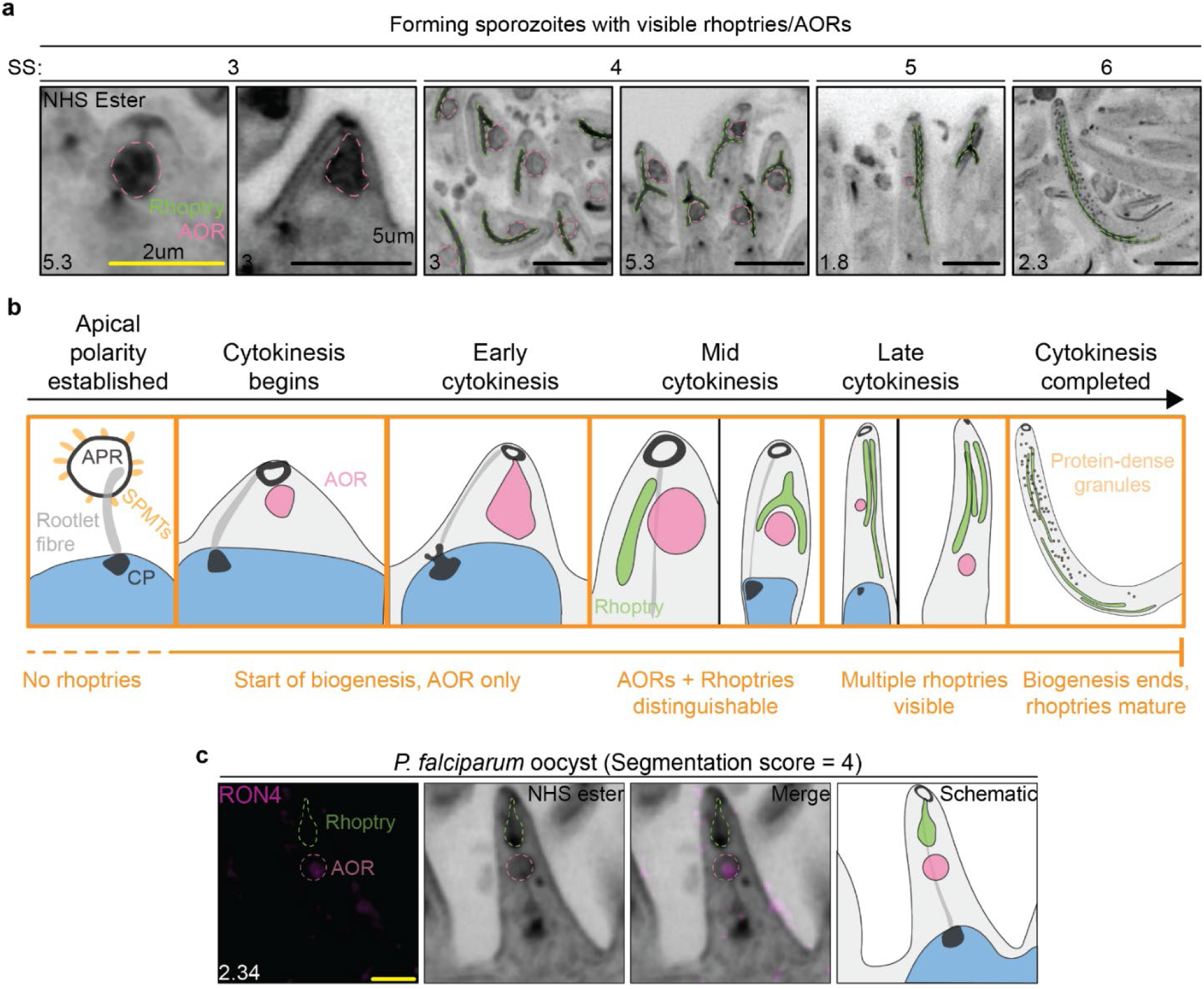
An inferred timeline for sporozoite rhoptry biogenesis. (**a**) Visualisation of either the rhoptries (green outline) or alternative or immature organelles distinguishable from mature rhoptry (AOR, pink outline) in *Plasmodium yoelii* sporozoites using NHS Ester (protein density, greyscale) at different stages of their development (SS = segmentation score). (**b**) Schematic for the inferred series of events in sporozoite rhoptry and AOR biogenesis with each cartoon representative of a forming sporozoite in the imaging dataset. Timeline also includes the formation of protein-dense granules in late sporozoite development. (**c**). Visualisation of a forming *P. falciparum* sporozoite (segmentation stage 4) stained with anti-RON4 (rhoptry neck) antibodies. Outlines show inferred positions of AOR and rhoptry. Scale bars: black = 5 µm, yellow = 2 µm. Number in bottom left corner = image z-depth in µm.

Rhoptries were not visible while oocysts were still undergoing mitosis, prior to sporozoite formation (segmentation score 1). This directly contrasts to blood-stage merozoites, where rhoptry duplication and segregation appears coupled to rounds of mitosis and nuclear division^25^. After the establishment of apical polarity and visualization of the apical polar ring (segmentation score 2-3), we observed a rhoptry-like structure that has previously been termed the AOR^14^ (Figure 4a; SS 3). The AOR was present prior to a visibly mature rhoptry, suggesting the AOR forms before the first rhoptry. Following the formation of the AOR, forming sporozoites contained both AORs and visibly recognizable rhoptries (Figure 4a; SS 4). AORs and rhoptries frequently seemed to be intimately associated with each other, with an AOR often appearing to directly dock to a rhoptry, or a rhoptry forking around an AOR (Figure 4a; SS 4). These observations suggest that the AOR may dock to rhoptries, potentially as a biogenesis vehicle as previously suggested^27^. AORs remain visible until late in sporozoite segmentation, and typically at this stage forming sporozoites only have 2-3 rhoptries, suggesting that rhoptry biogenesis is still occurring (Figure 4a; SS 5). By the end of cytokinesis, in fully mature sporozoites, AORs were no longer visible, only rhoptries (Figure 4a; SS 6). While our data cannot directly confirm that membrane fusion occurs between rhoptries and AORs, we wanted to determine if AORs carry rhoptry cargo. As we lacked access to a rhoptry-specific antibody for *P. yoelii*, we performed MoTissU-ExM on *P. falciparum*-infected midguts stained with antibodies against the rhoptry neck marker RON4. These images clearly showed that AORs carry rhoptry cargo (Figure 4c). Collectively, these observations suggest that sporozoite rhoptry biogenesis is uncoupled from mitosis/nuclear division, begins after the start of segmentation, ends just before the formation of fully mature sporozoites, and confirms the hypothesis that AORs carry rhoptry cargo.

In most oocysts imaged there were bundles of rhoptries and AORs in the centre of the sporoblast, not incorporated into any of the forming sporozoites (Figure S5). It is unclear whether these rhoptries and AORs are formed inside sporozoites and subsequently ejected, or if they are formed Golgi in the sporoblast. The central region of the sporoblast, which does not contain sporozoites, will eventually form the residual body upon oocyst egress and so these observations suggest that the oocyst residual body contains rhoptries.

#### Rhoptry necks and bulbs can be visually distinguished in *P. falciparum* sporozoites

Rhoptry biogenesis follows a specific sequence during blood-stage development: the rhoptry bulb forms first, followed by the rhoptry neck^21,25,26,38–40^. In *P. yoelii* and *P. berghei* sporozoites, however, the rhoptries were largely uniform in diameter with the bulb and neck not easily distinguishable from each other based on morphology alone (Figure 4). We were able to access and prepare a small number of *P. falciparum*-infected midguts by MoTissU-ExM, again staining with NHS Ester (Figure S6). By contrast to the other species we have imaged, rhoptries of forming *P. falciparum* sporozoites have discernable neck and bulb regions (Figure S6a). From early rhoptry biogenesis (segmentation score 3) until sporozoites are most of the way through cytokinesis (segmentation score 5) this rhoptry neck/bulb distinction remains clear (Figure S6b). By contrast, the distinction is not as easily discernable in rhoptries from mature oocyst sporozoites (segmentation score 6-7; Figure S6b). Except for the ability to distinguish rhoptry necks and bulbs, the process and timing of rhoptry biogenesis (Figure S6b), along with rhoptry number (Figure S6c) appear to be the same in *P. falciparum* as *P. yoelii* and *P. berghei*.

### Sporozoites use two rhoptries during salivary gland invasion

It is currently unknown how many rhoptries a sporozoite has, whether this number changes during SG invasion, or if there are rhoptries ‘specialised’ for each of the sporozoite invasion events. Previous reports in the literature suggest sporozoites have 2-6 rhoptries^8,22,41^, but it is unclear whether this range occurs between sporozoites, between lifecycle stages, or between *Plasmodium* species. Sporozoite rhoptries are long, slender, and flexible. Because of these properties, rhoptries cannot be individually visualised by conventional light microscopy, while thin-section transmission electron microscopy may either not sample a rhoptry in a particular slice or sample the same rhoptry multiple times; precluding accurate estimation of sporozoite rhoptry number. Using MoTissU-ExM, we were able to distinguish individual rhoptries. To determine rhoptry number, and whether this changes during SG invasion, we counted rhoptries from: mature oocyst sporozoites (segmentation score 6-7), haemolymph sporozoites, sporozoites inside SG epithelial cell, and secretory cavity sporozoites (Figure 5a). Mature oocyst sporozoites and haemolymph sporozoites both contained a median of 4 rhoptries (oocyst - range 3-5, ± 0.55 SD; haemolymph – range 2-6, ± 0.78 SD) (Figure 5a). By contrast, SG epithelial cell and secretory cavity sporozoites contained a median of 2 rhoptries (epithelial cell – range 1-3, ± 0.34 SD; secretory cavity – range 0-2, ± 0.56) (Figure 5a). The loss of rhoptries between the haemolymph and SG epithelial cell suggests that sporozoites use up two of their rhoptries during SG invasion. Additionally, it suggests that entry into the secretory cavity may be rhoptry independent. Of note, we were only able to obtain *P. berghei* haemolymph sporozoites, while the other three anatomical niches were examined using *P. yoelii*.

**Figure 5:**
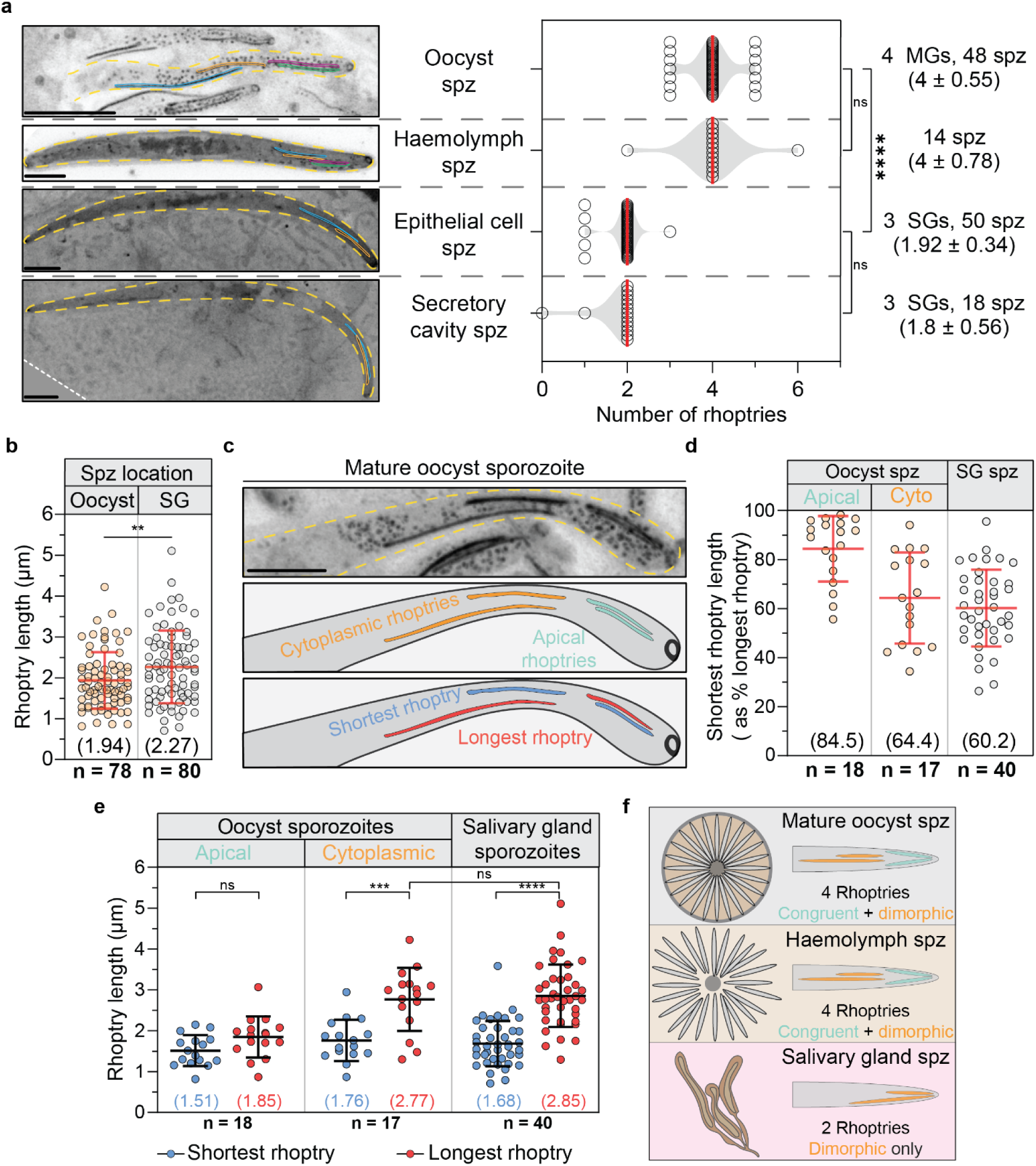
Sporozoites have morphologically distinguishable rhoptry pairs and use one during salivary gland invasion. (**a**) Based on protein density staining (NHS Ester, greyscale), rhoptry number was determined for mature oocyst sporozoites, haemolymph sporozoites, SG epithelial cell, and SG secretory cavity sporozoites. Secretory cavity sporozoite images was rotated for presentation purposes, dotted white line indicates border of original image. Dashed yellow line indicates sporozoite plasma membrane, each rhoptry outlined in a different colour. (**b**) Measurement of rhoptry length in mature oocyst sporozoites and SG sporozoites. (**c**) In mature oocyst sporozoites, rhoptry pairs were defined as either apical or cytoplasmic (cyto). (**d**-**e**) The length of these rhoptry pairs were measured both relative to each other (**d**), and individually (**e**), alongside the single rhoptry pair in SG sporozoites. (**f**) Schematic displaying the position of rhoptry pairs that were measured in (**d** and **e**). Based on the change in rhoptry number and morphological distinction, we propose to name the two oocyst sporozoite rhoptry pairs congruent (same length, previously apical) and dimorphic (differing in length, previously cytoplasmic), and suggest that following SG invasion only the dimorphic rhoptries remain. Scale bars = 5 µm. All lengths are corrected for expansion factor. Error bars = SD. ns = p > 0.05, ** = p < 0.01, *** = p <0.001, **** = p < 0.0001 by ANOVA (**a** and **e**) or two-tailed unpaired t-test (**b**).

### Sporozoites contain morphologically distinct rhoptry pairs

Sporozoites must invade both the mosquito SG and hepatocytes following transmission to their vertebrate host^34,42^. Considering the loss of rhoptries following SG invasion, we hypothesized that sporozoites may possess rhoptries that are specialized for each of these two invasion events, which might be reflected in their morphology.

To investigate this, we first measured rhoptry length on both mature oocyst sporozoites (segmentation score 6-7) and SG sporozoites (Figure 5b). We found a small, but significant difference in rhoptry length between oocyst and SG sporozoites (Figure 5b). While imaging mature oocyst sporozoites, we frequently observed two distinct pairs of rhoptries: one pair at the apical tip of the sporozoite (apical rhoptries), and another further recessed into the cytoplasm (cytoplasmic rhoptries) (Figure 5c). When comparing the length of these rhoptry pairs, we found that the apical rhoptries were of approximately equal length (Figures 5d and 5e) while for the cytoplasmic rhoptry pair, one rhoptry was consistently about ∼40% longer than the other (Figures 5d and 5e). Due to these differences, we will refer to the apical rhoptry pair as the ‘congruent rhoptries’ and the cytoplasmic rhoptry pair as the ‘dimorphic rhoptries’. When imaging SG sporozoites we typically found only a single rhoptry pair, present at the apical end of the sporozoite. When measured, the rhoptry pair from SG sporozoites were approximately equal in length to the dimorphic rhoptries from oocyst sporozoites (Figures 5d and 5e). This suggests that the rhoptry pair present at the apical end of SG sporozoites is likely the dimorphic pair from mature oocyst sporozoites. By extension, this suggests that the congruent rhoptries are used up during SG invasion, leaving the dimorphic rhoptries for hepatocyte invasion (Figure 5f). While we cannot yet confirm whether these different rhoptry pairs carry different cargo, these data show that rhoptry pairs can be morphologically distinguished and suggest that sporozoites may have rhoptry pairs specialized for each of their invasion events.

### Sporozoites not expressing RON11 have visually aberrant rhoptries

Having defined a timeline for rhoptry biogenesis and shown that a rhoptry pair is used during SG invasion, we wanted to use this information to help characterize the functions of a rhoptry protein. Previously, no proteins were known to be directly involved in sporozoite rhoptry biogenesis.

A recent study in *P. falciparum* used U-ExM to show that Rhoptry neck protein 11 (*Pf*RON11) was involved in rhoptry biogenesis, with P*f*RON11 knockdown leading to the formation of merozoites that contain only a single rhoptry and were defective for red blood cell invasion^26^. RON11 is a large protein, containing a signal peptide, dual-EF hand domains, and 7 transmembrane helices that span the rhoptry membrane (Figure S7). It is likely that the rhoptry luminal portion of RON11 has a function in merozoite invasion, while the cytosolically-exposed portion of RON11 coordinates its rhoptry biogenesis function. RON11 is expressed in *P. berghei* sporozoites and localizes to the rhoptries^14^. Additionally, the generation of a promoter swap mutant that drastically reduced *Pbron11*expression (*Pb*RON11^cKD^) led to the formation of sporozoites with substantially reduced ability to invade either the mosquito SG or hepatocytes^14^. Given the role of RON11 in *P. falciparum* merozoite rhoptry biogenesis, along with its localization and importance for sporozoite invasion, we sought to determine if RON11 is involved in sporozoite rhoptry biogenesis.

We obtained *P. berghei* RON11 promoter swap (*Pb*RON11^cKD^) and control (*Pb*RON11^ctrl^)^14^ parasites (a generous gift from Prof. Tomoko Ishino) (Figure 6a). Using *Pb*RON11^ctrl^ parasites, we observed that the segmentation score and timeline for rhoptry biogenesis in *P. berghei* were morphologically indistinguishable for what we had defined in *P. yoelii* (Figure 6b). Assessing rhoptry biogenesis in both *Pb*RON11^cKD^ parasites, we noticed no differences in early rhoptry biogenesis, with parasite forming rhoptries and AORs (Figures 6b and 6c). However, by mid-segmentation (segmentation score 4-5), many *Pb*RON11^cKD^ sporozoites had formed visibly aberrant rhoptries (Figure 6c). Typically, *Pb*RON11^ctrl^ rhoptries were slender and roughly uniform in diameter. By contrast, *Pb*RON11^cKD^ rhoptries frequently formed large protein-dense spheres (Figure 6c). It is currently unclear what leads to the formation of these aberrant rhoptries, however, they persisted until the formation of mature sporozoites (segmentation score 6-7). Within a single oocyst, typically all or none of the sporozoites exhibited these rhoptry aberrations, rather than a proportion of forming sporozoites in each oocyst. When quantified, visually aberrant rhoptries were observed in 23/42 (54.8%) *PbRON*11^cKD^ oocysts, and 0/26 (0%) *Pb*RON11^ctrl^ oocysts (Figure 6d). While these rhoptry defects were visually striking, it was previously shown that SG invasion of *RON*11^cKD^ sporozoites was reduced by >99%^14^. It is unlikely that approximately half of oocysts having visibly aberrant rhoptries fully explains this SG invasion defect.

**Figure 6:**
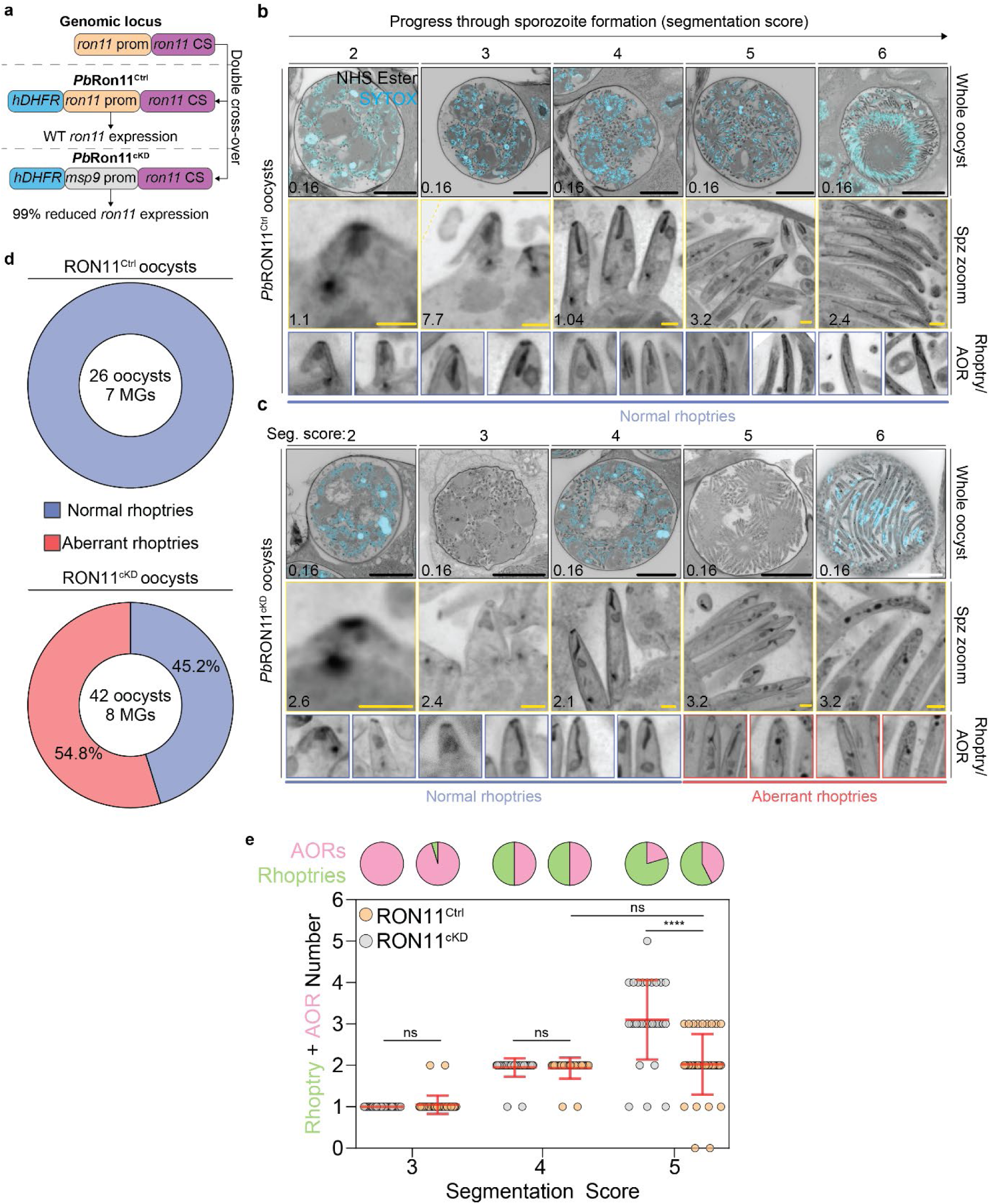
RON11^cKD^ sporozoites have visibly aberrant rhoptries. (**a**) Schematic showing Ron11^Ctrl^ and Ron11^cKD^ parasite lines as previously established^14^. Mosquito midguts were dissected with either *Plasmodium berghei* RON11^Ctrl^ (**b**) or RON11^cKD^ (**c**) parasites and prepared for MoTissU-ExM, stained with the protein density dye NHS Ester (greyscale) and SYTOX Deep Red (DNA, cyan). Representative images of oocysts at different stages of their sporozoite and rhoptry development are shown. Additional examples of ‘normal’ rhoptries are shown for RON11^ctrl^ and RON11^cKD^ with segmentation scores of 2-4, while examples of of ‘aberrant’ rhoptries are shown for RON11^cKD^ with segmentation scores of 5-6. Scale bars: black = 50 µm, white = 20 µm, yellow = 2 µm. (**d**) The presence of normal or ‘aberrant’ in RON11^Ctrl^ or RON11^cKD^ oocysts. (**e**) Quantification of the number of rhoptries or AORs in RON11^Ctrl^ or RON11^cKD^ forming sporozoites at different stages of segmentation. Pie charts represent the total proportion of AORs and rhoptries counted. ns = p > 0.05, **** = p <0.0001 by ANOVA. Error bars = SD.

### Sporozoites not expressing RON11 have half the expected number of rhoptries

While visualising *Pb*RON11^cKD^ and *Pb*RON11^ctrl^ parasites we observed that following appearance of aberrant rhoptries (segmentation stage 5 onwards), forming *Pb*RON11^cKD^ sporozoites contained fewer rhoptries and AORs than controls (Figure 6e). This suggested that rhoptry biogenesis may occur more slowly, or arrest earlier in *Pb*RON11^cKD^ parasites. Considering this observation, and the previous report that knockdown of *Pf*RON11 led to formation of merozoites with only a single rhoptry^26^, we wanted to determine the number of rhoptries in *Pb*RON11^cKD^ sporozoites. We had previously determined that mature oocyst sporozoites typically have 4 rhoptries, while SG sporozoites have only 2. For mature oocyst sporozoites, *Pb*RON11^ctrl^ parasites had a median of 4 rhoptries (range 2-5, ± 0.57 SD) (Figure 7a), as expected. By contrast, *Pb*RON11^cKD^ oocyst sporozoites contained a median of only 2 rhoptries (range 1-4, ± 0.52 SD) (Figure 7a). This halving in rhoptry number in *Pb*RON11^cKD^ sporozoites mimics the single-rhoptry merozoite phenotype following *Pf*RON11 knockdown^26^.

**Figure 7:**
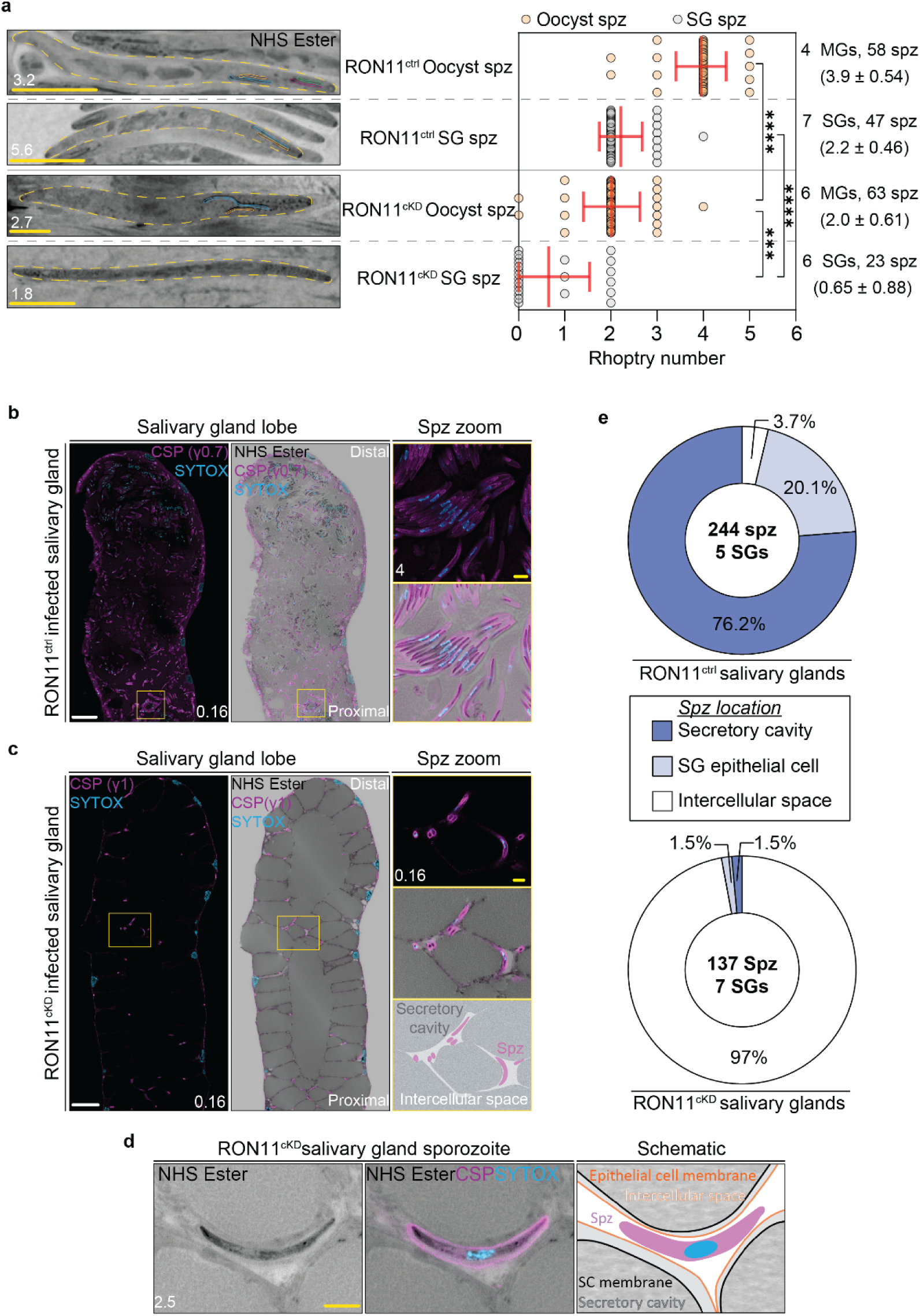
RON11^cKD^ sporozoites fail to invade the secretory cavity and have fewer rhoptries. (**a**) Quantification of rhoptry number in RON11^Ctrl^ and RON11^cKD^ parasites from both mature oocyst (MG) and SG sporozoites. Sporozoite membrane outlined in yellow, with each visible rhoptry outlined in a different colour. Error bars = SD. *** = p <0.001, **** = p <0.0001 by ANOVA. (**b**-**c**) SGs of mosquitoes infected with either *Plasmodium berghei* (**b**) RON11^Ctrl^ or (**c**) RON11^cKD^ parasites and prepared for MoTissU-ExM, stained with the protein density dye NHS Ester (greyscale), SYTOX Deep Red (DNA, cyan) and anti-CSP antibodies (sporozoite surface, magenta). The gamma (γ) for CSP fluorescence in (**b**) has been modified to 0.7 to better display spz at the centre of the SG. (**d**) Representative image of a RON11^cKD^ sporozoite (spz), which were often found inside the SG but not intracellular or inside the secretory cavity (SC). (**e**) Quantification of the location of sporozoites within the salivary gland. Scale bars: white = 100 µm, yellow = 10 µm

While previous studies showed that *Pb*RON11^cKD^ have a markedly reduced ability to invade the SG (>100-fold fewer SG sporozoites than *Pb*RON11^ctrl^)^14^, a small number of sporozoites were observed within SGs. Given the halving of mature oocyst sporozoite rhoptry number, and the previous observation that two rhoptries are used during SG invasion (Figure 5), we wanted to determine the number of rhoptries in *Pb*RON11^cKD^ SG sporozoites. As anticipated, *Pb*RON11^ctrl^ SG sporozoites contained a median of 2 rhoptries (range 2-4, ± 0.46 SD) (Figure 7a). Strikingly, *Pb*RON11^cKD^ SG sporozoites contained a median of 0 rhoptries (range 0-2, ± 0.88 SD) (Figure 7a). 64% of *Pb*RON11^cKD^ SG sporozoites contained no rhoptries at all, while 9% contained 1 rhoptry and 27% contained 2 rhoptries. The majority of *Pb*RON11^cKD^ mature oocyst sporozoites containing 2 rhoptries, while the majority of those in the SG do not contain any likely indicates that most sporozoites that make it to the SG still attempt to secrete their remaining rhoptries.

### Sporozoites lacking RON11 can cross the salivary gland basal lamina but fail to invade the epithelial cells

While imaging *Pb*RON11^cKD^ infected SGs, we noticed a peculiar sporozoite distribution. *Pb*RON11^ctrl^ sporozoites seemed to be relatively uniformly distributed throughout the SGs, invading the epithelial cells and secretory cavities of both the lateral and medial SG lobes (Figures 7b, S8a and S8b). By contrast, *Pb*RON11^cKD^ sporozoites were only very infrequently in either the secretory cavity or the SG epithelial cell (Figures 7c, S9a and S9b). *Pb*RON11^cKD^ sporozoites seemed mostly to be either between the basal lamina and SG epithelial cell or in a small cavity between epithelial cells, a region we collectively refer to as the ‘intercellular space’ (Figures 7c, 7d, and S9c). *Pb*RON11^cKD^ sporozoites in the intercellular space are best shown ‘slice-by-slice’ as in Supplementary Videos 1-8.

It has been shown that the presence of sporozoites in the secretory cavity disrupts the usual homogenous distribution of saliva and produces granular pattern instead^43^. In *Pb*RON11^ctrl^ infected SGs we almost always observed this effect, but not in *Pb*RON11^ctrl^ infected SGs (Figures 7b, 7c, S8a, and S9a). To determine if this was due to the absence of sporozoites in the secretory cavity, we screened SGs infected with *Pb*RON11^cKD^ or *Pb*RON11^ctrl^ parasites assessing if the SG was infected and then if the secretory cavity had been invaded (Figures S8c and S9d). A larger proportion of *Pb*RON11^cKD^ (28%) SGs were uninfected compared to *Pb*RON11^ctrl^ (10%) (Figures S8c and S9d) but mosquitoes were not screened for infection prior to SG dissection. In *Pb*RON11^ctrl^ infected SGs, 100% (35/35) contained sporozoites in the secretory cavity (Figure S8c). Conversely only 24% (14/59) of *Pb*RON11^cKD^ infected SGs contained sporozoites within the secretory cavity (Figure S9d). In a subset of infected SGs, we further refined the location of sporozoites. In *Pb*RON11^ctrl^-infected SGs, the majority of sporozoites were within the secretory cavity (76.2%), with the SG epithelial cell containing a further 20.1% of the sporozoites and the intercellular space containing only 3.7% of sporozoites (Figure 7e). By contrast, in *Pb*RON11^cKD^-infected SGs 97% of sporozoites were in the intercellular space, with only 1.5% of sporozoites in either the SG epithelial cell or secretory cavity (Figure 7e). Collectively this shows that RON11 is important for invasion of the SG epithelial cell. Sporozoites unable to enter SG epithelial cell would subsequently be unable to enter the secretory cavity and therefore would be unable to be transmitted during blood-feeding; explaining why *Pb*RON11^cKD^ parasites are not transmissible^14^.

## DISCUSSION

### Summary

In this study we illustrate the utility of MoTissU-ExM for studying the biology of *Plasmodium* oocysts and SG sporozoites. We visualize the intranuclear microtubule structures coordinating oocyst mitosis, the oocyst/sporozoite basal complex, apical polar ring, and centriolar plaque; all of which have not previously been observed using light microscopy. Given the lack of synchronicity in sporozoite formation between oocysts, we then established the segmentation score as a proxy for sporozoite development, against which the relative timing of other cellular events can be compared. Using this segmentation score, we then developed a timeline for the biogenesis of rhoptries in forming oocyst sporozoites. *Pb*RON11^ctrl^-infected SGs. Using *Pb*RON11^cKD^ parasites and the established timeline for rhoptry biogenesis, we identify when RON11 is involved in rhoptry biogenesis. Further, we show that *Pb*RON11^cKD^ parasites have a reduced number of rhoptries and that while they could cross the SG basal lamina, they are unable to enter the SG epithelial cell or secretory cavity where they need to reach to enable transmission.

### Progress through cytokinesis as a proxy for sporozoite development

When imaging fixed cells temporal dynamics have to be inferred from other experimental cues, typically either absolute time or by using a biological clock. Currently, there is no way to synchronise when ookinete invasion occurs and so the development of each oocyst within a single midgut is asynchronous. Considering this, a biological clock must be used to compare events happening in different oocysts. We had previously used cytokinesis, or segmentation, as a biological clock in blood-stage malaria parasites^25^ and so established the segmentation score to do the same in oocysts. This enabled us to define a timeline for rhoptry biogenesis, and we hope will be utilised by other research groups to characterize the temporal dynamics of their processes of interest. Currently, the biggest drawback of the segmentation score is that it cannot be used in oocysts that have not yet begun cytokinesis. Considering that oocyst size does not plateau until cytokinesis beings^4^, measuring oocyst diameter may represent a useful biological clock in these cases.

### Comparison between Plasmodium spp

To date, the only well documented differences in sporozoite ultrastructure between *Plasmodium* spp. is their length and number of subpellicular microtubules^4,44,45^. Despite our extensive analysis of both forming sporozoites in oocysts, and SG sporozoites, this is essentially still the case. During the biogenesis of *P. falciparum* sporozoite rhoptries, their neck and bulb regions could be visibly discerned, unlike *P. berghei* or *P. yoelii*, but in mature oocyst sporozoites this distinction was lost. With this sole exception, all other cellular structures imaged across *P. falciparum*, *P. yoelii*, and *P. berghei*, were visibly indistinguishable. Considering that these species of malaria parasites represent up to 100 million years of evolutionary divergence^46^ (noting the caveat of difficulty in estimating *Plasmodium* evolutionary divergence^47^), this exhibits a remarkable evolutionary conservation of sporozoite ultrastructure.

One unexpected finding of this study was the presence of a large number of rhoptries, AORs and nuclei in the region of the sporoblast that doesn’t get incorporated into any of the forming sporozoites. As this structure remains in the oocyst following sporozoite egress, it represents the oocyst residual body. While this structure has been visualized previously using electron microscopy^24^, to our knowledge its contents have not been investigated. Investigation of the contents of residual bodies in *Plasmodium* has been limited, but recent studies have shown that it can contain mitochondria and rarely apicoplast^25,48^. By contrast, the *Toxoplasma* tachyzoite residual body is more well studied and has been shown to contain nuclei, rhoptries, micronemes, golgi, dense granules, and mitochondria^49–51^.

### Sporozoite rhoptry biogenesis is dissimilar from blood-stage merozoite rhoptry biogenesis

During rhoptry biogenesis of blood-stage parasites, the presence, duplication, and inheritance of rhoptries into daughter merozoites are intrinsically linked to mitosis and nuclear division^25^. In short, during each round of mitosis the merozoite rhoptries duplicate alongside the CP and are inherited with each of the sister nuclei following nuclear division^25^. From our observations, it seems that sporozoite rhoptry biogenesis begins around the same time that oocysts stop undergoing mitosis. While sporozoite rhoptry biogenesis seems to lack the association with mitosis observed in merozoites, it does appear that the timing of rhoptry biogenesis in both forming merozoites and sporozoites occurs similarly with respect to cytokinesis (segmentation). In both instances, the ability to visualise rhoptries by protein density coincides with the ability to see the basal complex; the first indication that cytokinesis has begun^25^. Curiously in *Toxoplasma gondii*, *de novo* rhoptry biogenesis also begins immediately after the start of cytokinesis^51^, suggesting that an association between the onset of cytokinesis and rhoptry biogenesis may be shared across Apicomplexa. Like *Plasmodium* sporozoites, rhoptry biogenesis in both *T. gondii*^51^ and *Cryptosporidium*^52^ appears to be uncoupled from mitosis and nuclear division, suggesting that this association may be a unique adaptation of *Plasmodium* merozoite rhoptry biogenesis.

### Salivary gland invasion requires rhoptries

Until relatively recently, it was unclear if rhoptries were required for sporozoite invasion of the SG^5^, with suggestions that this process was akin to traversal^53,54^ rather than productive invasion^55^. Since 2019, however, a range of rhoptry proteins have been shown to be involved in SG invasion^13–15^, and a 2022 study provided the first direct evidence that SG invasion involves the formation of a canonical tight junction^12^. Following invasion events by *Plasmodium* merozoites, the rhoptry pair that facilitate invasion are ‘used up’ and disassembled rapidly following invasion^25,56^. We show that this loss of a rhoptry pair occurs following sporozoite invasion of the SG epithelial cell but not the secretory cavity. This finding strengthens previous observations that SG invasion involves formation of a tight-junction and rhoptry secretion^12^, and therefore represents a ‘true invasion’ event and not cell traversal.

### Sporozoites may contain rhoptry pairs specialized for different invasion events

One of the most significant findings of this study is that oocyst sporozoites possess morphologically distinguishable pairs of rhoptries, termed congruent and dimorphic, and that following SG invasion, only the dimorphic rhoptries remain. Our interpretation of these observations is that the congruent rhoptries are used during SG invasion, and by extension that the dimorphic rhoptries would be used for hepatocyte invasion, where rhoptry involvement is known^23,57^. It is therefore logical to speculate that the two pairs of rhoptries may be specialized for different invasion events; congruent for the SG, dimorphic for hepatocytes. Considering the cell biological and evolutionary differences between a mosquito SG epithelial cell and a hepatocyte, it would be unsurprising if at least a small subset of different proteins are required to facilitate invasion in these two cell types, however no functional data exists to support this hypothesis. Despite our observation of morphological differences in rhoptry pairs, our data does not provide any evidence that these rhoptry pairs are functionally different or carry different cargo. The phenotype of the knockdown of RON11 does not aid in this interpretation as there does not seem to be a consistent trend as to the rhoptry pair that are missing in RON11^cKD^ oocyst sporozoites. Additionally, it has previously been shown that the portion of RON11 present in the rhoptry lumen plays an important role in merozoite invasion that is unrelated to the function of RON11 in rhoptry biogenesis^26^. Considering this, it is likely that the RON11^cKD^ parasites used in this study have combinatorial defects that effect both rhoptry biogenesis and invasion, subsequently inhibiting both SG invasion and hepatocyte invasion^14^.

### Presence of PbRON11^cKD^ sporozoites in the SG intercellular space

The majority of PbRON11^cKD^-infected mosquitoes contained some sporozoites in their SGs, but these sporozoites were only rarely inside either the SG epithelial cell or secretory cavity. Instead, sporozoites were either between the basal lamina and SG epithelial cells, or in the intercellular space between SG epithelial cells. Observation of sporozoites in cavities between SG epithelial cells was unexpected as they typically form very narrow tight junctions that are difficult to discern even by electron microscopy^43^. To the best of our knowledge, the ability of sporozoites to cross the basal lamina and accumulate in the SG intercellular space has never previously been reported. Based on these observations, we hypothesise that the sporozoites that fail SG epithelial cell invasion are capable of forming these cavities between SG epithelial cells, although it is currently unclear how this could be experimentally determined.

Despite their ability to cross the basal lamina, PbRON11^cKD^-infected SGs contain <1% the number of sporozoites as control SGs despite similar numbers of haemolymph sporozoites^14^. It is unclear if this represents an additional defect in the ability of these sporozoites to cross the basal lamina, or if this confined space can only physically accommodate a small number of sporozoites. It has been shown that PbRON11^cKD^ have a motility defect^14^ and so this may be associated with a defect in basal lamina traversal. It has previously been shown that infection of SG epithelial cells and subsequently the secretory cavity by large numbers of sporozoites damages the SG^43^. It is possible that this damage makes the SG more permissive to further sporozoite invasion. If this is the case, the drastic reduction in secretory cavity invasion of PbRON11^cKD^ sporozoites may be directly responsible its overall reduced number of SG sporozoites.

### Future applications

Many of the insights in this study would have been difficult or impossible to attain without the use of expansion microscopy. It is therefore extraordinarily exciting to consider the new biology to be uncovered using this technique looking to the future. In terms of *Plasmodium* biology, we suspect MoTissU-ExM can be adapted to investigate migration and traversal by ookinetes followed by early oocyst formation *in situ*. Additionally, MoTissU-ExM enhances our ability to study parasite-mosquito interactions, by enabling visualization of mosquito tissue architecture and parasite ultrastructure in the same sample. Utilising the segmentation score we have developed in this study, MoTissU-ExM may represent a powerful tool to assess the effect of antiparasitic drugs or insecticides on sporozoite development.

In other tissue types, expansion microscopy has been multiplexed with spatial transcriptomics and proteomics^58–60^, opening the possibility to perform these techniques on infected mosquito tissues. More broadly, a myriad of pathogens of humans, animals, and plants are ingested into the midgut of an arthropod and translocate to the SG to enable their transmission. Therefore, MoTissU-ExM can likely be adapted to study a multitude of arthropods including sandflies, ticks, hoppers, or other genera of mosquitoes and may enable new understanding into the biology of diseases transmitted by these arthropods such as leishmaniasis, theileriosis, bacterial leaf scorch (*Xylella*), dog heart worm (*Dirofilaria*), or Zika.

## MATERIALS AND METHODS

### Mosquito rearing, infections, and dissection

#### Mosquitoes

Experiments with *Plasmodium falciparum* were performed using *Anopheles gambiae* Keele Strain^61^, or *Anopheles stephensi* (Nijmegen)^62^. Experiments with *Plasmodium berghei* were performed using *Anopheles gambiae* Keele strain. Experiments with *Plasmodium yoelii* were performed using *Anopheles stephensi* (Seattle Children’s Research Institute).

#### Mosquito rearing

All mosquitoes were reared at 27 °C and 80% humidity under a 12-h light/dark cycle under standard laboratory conditions and maintained with 10% sugar syrup solution during adult stages. After blood-feeding *P. berghei*-infected mosquitoes were kept at 19 °C, while *P. falciparum* were kept at 25 °C.

#### Plasmodium yoelii infection

For *P. yoelii* infection, female 6–8-week-old Swiss Webster mice (Harlan, Indianapolis, IN) were injected with blood stage *P. yoelii* (17XNL) parasites to begin the growth cycle. Mouse handling was conducted in accordance with Settle Children’s Research Institute Animal Care and Use Committee-approved protocols. *Anopheles stephensi* mosquitoes were allowed to feed on infected mice after gametocyte exflagellation was observed.

#### *Plasmodium berghei* infection

For infection, female *Anopheles gambiae* Keele strain mosquitoes, aged 4–5 days, were allowed to feed on *P. berghei*-infected mice with an exflagellation rate of 1 exflagellants per 40 × microscopic field. Midguts were dissected 10 days post infection and salivary glands 21 days.

#### *Plasmodium falciparum* infection

Female *Anopheles gambiae* Keele strain, or *Anopheles stephensi* (Nijmegen), mosquitoes were infected with *P. falciparum* NF54 via membrane feeding using reconstituted human blood^63^. The asexual blood-stage cultures were maintained in vitro using O+ erythrocytes at a 4% hematocrit, as previously described^63^. For mosquito infections, a suspension of human RBCs was mixed with human serum (Interstate blood bank) at 50% hematocrit and diluted to 0.05% gametocytemia before being fed to the mosquitoes. Midguts were dissected 10 days post infection.

#### Haemolymph sporozoites

Hemolymph was collected by perfusion from *berghei*-infected mosquitoes. Mosquitoes were anesthetized on ice and placed individually under a stereoscope on parafilm. An incision was made on the lateral region of the VI-VII abdominal segments using a U-100 insulin syringe. A heat-stretched capillary needle filled with sterile PBS was inserted laterally into the thorax, and approximately 10 μL of sterile PBS was injected to displace the hemolymph. The hemolymph flowed through the abdominal incision and was collected with a pipette (P20, Gilson) before settling on 12 mm coverslips for ∼1 hr and fixation in 4% paraformaldehyde.

#### Salivary gland dissection

Salivary glands were dissected from either *P. yoelii* (14 dpi) or *P. berghei*-infected (21 dpi) mosquitoes. Following dissection, SGs were immediately fixed in 4% paraformaldehyde for 30 minutes at room temperature followed by 150 minutes at 4 °C and being washed and stored in PBS.

#### Midgut dissection

Mosquito midguts were dissected from either *P. yoelii*, *P. berghei*, or *P. falciparum*-infected mosquitoes and placed in PBS at room temperature. Midguts were subsequently immersed in 4% paraformaldehyde for 30 minutes at room temperature followed by 150 minutes at 4 °C and being washed and stored in PBS.

#### Mouse details and mouse handling

For experiments relating to *P. berghei,* all animal procedures were performed in strict accordance with the National Institutes of Health (NIH) guidelines under protocols approved by the National Institute of Allergy and Infectious Diseases Animal Care and Use Committees (NIAID ACUC). The studies were done following approved animal study proposals LMVR-22.

All mice used in this study were Swiss Webster mice purchased from Charles River Laboratories.

#### Salivary gland sporozoite isolation

SG sporozoites were isolated from hand-dissected salivary glands at day 14 or 15 post-blood meal. Pooled salivary glands were mechanically ground using a pestle in 3 sets of 30 grinds, followed by centrifugation for 2 min at 60 ×*g*. Supernatant containing sporozoites was collected and sporozoites were activated with 20% v/v FBS. Sporozoites were then pelleted by centrifugation at 1,500 ×*g* for 2 minutes and resuspended in 4% v/v paraformaldehyde.

##### Parasite strains

###### Plasmodium berghei

All *P. berghei* experiments in this study used strain ANKA were genetically modified to express a green fluorescent protein (GFP) driven by the elongation factor 1A (ef1α) promoter^64^. Additionally, the promoter locus of *ron11* had been modified to either encode the endogenous *ron11* promoter (RON11^Ctrl^), or the promoter of msp9 (RON11^cKD^), both with the addition of the hDHFR selectable marker^14^.

###### Plasmodium falciparum

All *P. falciparum* experiments in this study used strain NF54^65^ that had not been genetically modified.

###### Plasmodium yoelii

All *P. yoelii* experiments in this study used strain 17XNL (BEI Resources) that had not been genetically modified.

### Ultrastructure Expansion Microscopy (U-ExM)

All expansion microscopy in this study was performed as described in previous studies detailed below. All expansion microscopy experiments are fundamentally derivatives of the first U-ExM protocol^17,18^, with subsequent modifications of the protocol related to its use in *Plasmodium*^25,28,30^.

Briefly, samples were fixed in 4% paraformaldehyde and adhered to a 12 mm round coverslip coated with 0.1 mg/mL Poly-D-Lysine. After adherence, samples were incubated with anchoring solution (formaldehyde/acrylamide) overnight at 37 °C. After anchoring, coverslips were washed with 1x PBS and placed onto a piece of parafilm in a humidity chamber that had been cooled on ice. Activated monomer solution was made by mixing tetraethylmethylenediame (TEMED) and ammonium persulphate with monomer solution (sodium acrylate, bis-acrylamide, acrylamide, in PBS). A spot of activated monomer solution was placed on the parafilm, with the coverslip placed on top of this solution. Gels were then allowed to polymerise at 37 °C for 30 minutes. Polymerised gels were transferred to the wells of a 6-well plate containing 2 mL denaturation buffer (Sodium dodecyl sulphate, Tris, Sodium chloride in deionized water) and placed on an orbital shaker for 15 minutes to separate the gel and coverslip. Separated gels were then placed into a 1.5 mL Eppendorf tube containing denaturation buffer and capped before incubation at 95 °C in a heat block for 90 minutes. Following denaturation, gels were expanded through three 30 minute washes in deionized water. After initial expansion, gels were measured to determine expansion factor and either processed for antibody staining, or washed with 50% glycerol in water for cryopreservation. Stained gels were shrunken by washing twice for 15 minutes in 1x PBS before blocking in 3% bovine serum albumin in PBS (blocking buffer) for 30 minutes. After blocking, gels were incubated with primary antibodies, diluted in blocking buffer, overnight at room temperature on an orbital shaker. Following primary antibody incubation, gels were washed three times with PBS-Tween-20 (0.5% v/v; PBS-T) for ten minutes. After washing, gels were incubated for 2.5 hours in the dark with secondary antibody/fluorescent dye solution prepared in 1x PBS. After secondary antibody staining, gels were washed again three times with PBS-T before three rounds of expansion in deionized water for 30 minutes each. Gels stained with BODIPY TR or FL ceramide were incubated with the fluorophore overnight in water.

#### Isolated sporozoite U-ExM

Expansion microscopy of isolated sporozoites was performed as previously described^19^. Briefly, isolated SG or haemolymph sporozoites were allowed to settle onto a round 12 mm poly-D-lysine coated coverslip for 1 hr before fixation in 4% paraformaldehyde. Subsequently, these samples were processed as per the previously described U-ExM protocol.

### Mosquito Tissue Ultrastructure Expansion Microscopy (MoTissU-ExM)

MoTissU-ExM was developed to enable this study, but the development of this technique has previously been described in detail^19^. Briefly, isolated SGs or midguts were fixed in 4% paraformaldehyde. Following fixation, fixative was removed from the tissues and replaced with anchoring solution (formaldehyde/acrylamide), before anchoring overnight at 37 °C. After anchoring, tissues (typically 5-6 per coverslip) were transferred to a poly-D-lysine coated round 12 mm coverslip and concentrated by removing the anchoring solution. Gelation of tissues was then performed as described in the U-ExM protocol. Following gelation and gel polymerization, gels containing mosquito tissues were placed in 2 mL of denaturation buffer for 15 minutes on an orbital shaker to separate the gel and coverslip. After gel separation, individual SGs or midguts were cut out of the gel for individual staining and processing as described in the U-ExM protocol.

### Screening for salivary gland and secretory cavity invasion

SGs from either *Pb*RON11^cKD^ or *Pb*RON11^Ctrl^ infected mosquitoes were prepared by MoTissU-ExM as previously described, and stained with antibodies against *Pb*Circumsporozoite protein (CSP 3D11; BEI Resources^6^) to enable sporozoite visualization. SGs were first screened for infection on a low-magnification objective lens. Infected SGs were then screened on higher magnification objective lens (40 × C-apochromat autocorr M27 (water, 1.2 NA)), which enabled discernment of if sporozoites were present in the SG epithelial cell cytoplasm or secretory cavity.

Infected SGs lobes were first assessed to see whether their secretory cavities contained any sporozoites at all (Figures S8c and S9c). In Figure 7e, where this was quantified, presence of any sporozoites in the secretory cavity at all marked that SG as having sporozoites in the secretory cavity; without respect to sporozoite number. For some SGs, the entire volume of each of the three lobes could not be visualized using the high-magnification objective. For these SGs, the presence or absence of sporozoites in the imageable area of the secretory cavity (usually the vast majority) was interpreted as being representative of the presence or absence of sporozoites in the secretory cavity overall.

From a subset of these, tiled images of a single z-slice as close as possible to the central cavity of the lobe were taken. From these images, the proportion of sporozoites in the intercellular space, SG epithelial cell, and secretory cavity was quantified (Figure 7e). This quantification could not be meaningfully blinded because of the ∼90-fold change in sporozoite number between *Pb*RON11^cKD^ and *Pb*RON11^Ctrl^ infected SGs. Up to 50 sporozoites were counted in each SG image to not overrepresent highly-infected SGs. All images of *Pb*RON11^cKD^ infected salivary glands contained fewer than 50 sporozoites and so all sporozoites in the image were counted. All images of *Pb*RON11^Ctrl^ contained greater than 50 sporozoites, so to randomly sample 50 sporozoites the first 50 in the image from the distal most portion of the SG were quantified.

### Image acquisition

All images in this study were acquired using an LSM900 AxioObserver with Airyscan 2 (Zeiss, Oberkochen, Germany). Images were acquired on either a 5 × Fluar (air, 0.25 NA), 20 × Plan-apochromat (air, 0.8 NA), 40 × C-apochromat autocorr M27 (water, 1.2 NA), or 63 × Plan-apochromat (oil, 1.4 NA).

Tiled images of midguts were acquired using a 5 × Fluar (air, 0.25 NA) or 20 × Plan-apochromat (air, 0.8 NA) objective. Tiled images of SGs were acquired using a 40 × C-apochromat autocorr M27 (water, 1.2 NA) objective. All tiled images were made using the lowest possible zoom for that objective (typically 0.45x).

### Image processing

All images in this study underwent Airyscan processing using ‘moderate’ filter strength. Images acquired using the 5 × and 20 × objectives underwent 2D Airyscan processing, while images acquired using either the 40 × or 63 × objectives underwent 3D Airyscan processing. For images that contain NHS Ester, the gamma value of this channel was set to 0.45, rather than 1 for greater discernment of subcellular structures; as has previously been described^25^.

All images in this study were prepared and processed in, then exported from ZEN Blue Version 3.8 (Zeiss, Oberkochen, Germany). Tiled images were stitched using the ZEN Blue stitching function. The NHS Ester channel was always set as the reference channel, with all other channels adjusted relative to that. Tiled images used a 10% overlap and stitching was performed with both the ‘fuse tiles’ and ‘correct shading’ settings enabled.

### Image analysis

All reported distances were measured using the line function in ZEN either in 2D or 3D.

Where oocyst diameters were measured, these values were obtained by taking the maximum diameter of any oocyst on a single z-slice (2D measurement).

### Rhoptry measurement

Where rhoptry length was measured, these values were obtained by marking either end of the rhoptry and determining the distance between these two points using the 3D measurement function. For rhoptries that were not approximately a straight line in three-dimensions, additional points were added so the line always passed through the rhoptry, with the sum of these lines representing rhoptry length.

When performing the analysis of rhoptry pairs (Figures 5d and 5e) only oocyst sporozoites with 4 rhoptries (>75% of oocyst sporozoites), and SG sporozoites with 2 rhoptries (∼90% of SG sporozoites) were analysed. For oocyst sporozoites, the two rhoptries present closest to the apical polar ring were designated as the ‘apical pair’, while the two located more basally were designated the ‘cytoplasmic pair’. For all pairs, rhoptries were measured, with the shorter of these two designated the ‘shortest rhoptry’ and the other the ‘longest rhoptry’.

### Statistical analysis

All graphs presented in this study were generated using GraphPad PRISM (version 10). All error bars represent standard deviation. Where distances have been reported, these values have been corrected for the expansion factor (by dividing the measured value by 4.25) as previously described^19,25^.

### Stains and antibodies

A full list of stains, dyes, and antibodies used in this study can be found in Supplementary Tables 1 and 2.

### Protein structure and topology prediction

The structure of *Pb*RON11 (PBANKA_1327100) was predicted using the software Chai-1^66^. To make this prediction the *Pb*RON11 sequence was obtained from PlasmoDB (Release 68)^67,68^ and the amino acids 1-21 were removed due to their prediction as a signal peptide using Signal-P (Version 6.0)^69^ leaving aa22-955 for the Chai-1 prediction. TOPCONS was used to predict the presence of transmembrane helices and their topology in *Pb*RON11^70^. Residues with an annotation of ‘inside’ were interpreted to be present in the cytoplasm, while residues with an annotation of ‘outside’ were interpreted to be present in the rhoptry lumen. EF-Hand domain location was inferred from InterProScan^71^. *Pb*RON11 membrane topology map was constructed using Protter (Version 1.0)^72^. Predicted protein structure was visualized using the Mol* Viewer application^73^ available on Protein Data Bank^74^. Predicted structure was orientated to show the clearest delineation of transmembrane helices, rhoptry luminal loops and cytoplasmic loops.

### Data accessibility

All imaging data will be made publicly available following peer review, with confirmed completion of the imaging dataset. The authors intend to deposit and curate data through the database Dryad. Prior to peer review, the authors are happy to share all imaging data, which will be made available upon request to the corresponding authors.

## Supporting information

NIH Publishing Agreement & Manuscript Cover Sheet

## ACKNOWLEDGEMENTS

We greatly appreciate the financial support from the Indiana University School of Medicine Department of Pharmacology and Toxicology Postdoctoral Career Development Award that funded travel of BL to the lab of JVR. We acknowledge Professor Freddy Frischknecht (Universitätsklinikum Heidelberg) for insightful discussions on sporozoite biology, sharing of unpublished data, and critical reading of the manuscript. We thank Professor Tomoko Ishino (Institute of Science Tokyo) for provision of the RON11^cKD^ and RON11^Ctrl^ parasite lines. We acknowledge Nedal Darif (EMBL, Universität Heidelberg) and Meghan Zadow (University of Adelaide) for critical reading of the manuscript.

## AUTHOR CONTRIBUTIONS

**Benjamin Liffner** – Conceptualization, Data curation, Formal analysis, Funding acquisition, Investigation, Project administration, Supervision, Visualization, Writing – original draft, Writing – review & editing

**Thiago Luiz Alves e Silva** – Investigation, Methodology, Writing – review & editing

**Elizabeth Glennon** – Investigation, Writing – review & editing

**Veronica Primavera** – Investigation, Writing – review & editing

**Elaine Hoffman** - Investigation

**Alexis Kaushansky** – Conceptualization, Resources, Supervision, Writing – review & editing

**Joel-Vega Rodriguez** – Conceptualization, Resources, Supervision, Writing – review & editing

**Sabrina Absalon** – Conceptualization, Data curation, Formal analysis, Funding acquisition, Investigation, Resources, Project administration, Supervision, Writing – review & editing

