## Supplementary Information for "Unlocking new understanding of *Plasmodium* sporozoite biology with expansion microscopy"

**Supplementary Table 1:** **Summary of all antibodies used in this study.**

| Antibody (Ab) | Ab species | Ab source | Ab concentration | Reference/cat no. |
|---|---|---|---|---|
| Anti-*Pb*circumsporozoite protein (3D11) | Mouse | BEI Resources | 1: 1000 | MRA-100A[1] |
| Anti-mouse IgG Alexa Fluor 555 (H+L) | Goat | Invitrogen | 1:500 (2mg/ml stock) | A21422 |
| Anti-mouse IgG Alexa Fluor 488 Superclonal (H+L) | Goat | Invitrogen | 1:500 (1mg/ml) | A28175 |
| Anti-alpha tubulin (Clone B-5-1-2) | Mouse (IgG1) | ThermoFisher | 1:500 | 32-2500 |
| Anti-polyE (IN105) | Rabbit | Adipogen | 1:500 | AG-25B-0030-C050 |
| Anti-BIP | Rabbit | Gift from Dvorin Lab | 1:2000 | [2] |
| Anti-ERD2 (MRA-1) | Rabbit | BEI Resources MR4 | 1:2000 | [3] |
| Anti-RON4 | Mouse | Gift from Arnab Pain | 1:200 | [4] |
| Anti-*Pv*Circumsporozoite protein (2F2) | Mouse | | 1:1000 | [5] |
| Anti-Rabbit IgG Alexa Fluor 488 | Goat | ThermoFisher | 1:500 | A11034 |
| Anti-Rabbit IgG Alexa Fluor 555 | Goat | ThermoFisher | 1:500 | A21428 |

**Supplementary Table 2:** **Summary of all fluorescent dyes and stains used in this study.**

| Stain/dye | Source | Concentration | Step of protocol applied | Cat no. |
|---|---|---|---|---|
| NHS Ester Alexa Fluor 405 | ThermoFisher | 1:250 (2 mg/mL stock in DMSO) | Secondary Ab | A30000 |
| BODIPY-TR-Ceramide | ThermoFisher | 1:500 (1 mM stock in DMSO) | Post expansion | D7540 |
| BODIPY-FL-$C_5$-Ceramide | ThermoFisher | 1:500 (1 mM stock in DMSO) | Post expansion | D3521 |
| SYTOX Deep Red | ThermoFisher | 1:1000 (1 mM stock in DMSO) | Secondary Ab | S11381 |

**Supplementary Table 3: Summary of reagents and materials used in this study.**

| Reagent/material name | Reagent supplier | Catalogue number |
|---|---|---|
| Poly-D-Lysine 0.1mg/mL solution | Gibco | A3890401 |
| Ammonium persulfate (APS) powder | ThermoFisher | 17874 |
| Tetramethylethylenediamine (TEMED) | ThermoFisher | 17919 |
| Formaldehyde 36.5-38% solution | Sigma | 8775 |
| Acrylamide 40% solution | Sigma | A4058 |
| N,N'-methylenebisacrylamide (BIS) 2% | Sigma | M1533 |
| Sodium acrylate >97% powder | ThermoFisher | 408220 |
| Propyl gallate 98% powder | ThermoFisher | 131581000 |
| TRIS Base | RPI | T60040-250.0 |
| Sodium dodecyl sulfate micro-pellets | RPI | L22040-500.0 |
| Sodium chloride | RPI | S23020-1000.0 |
| Glycerol | Fisher | BP229-4 |
| 10x PBS | Sigma | 806552 |
| TWEEN-20 | Sigma | P1379 |
| Smart plastic razors | Sigma | Z740503 |
| 12 mm round coverslips | Fisher | NC1129240 |
| 35 mm Cellvis #1.5 glass bottomed dishes | Fisher | NC0409658 |
| Parafilm | Sigma | P7793 |
| Dissecting forceps | Fisher | NC9889584 |

**Supplementary Table 4: Summary of all solutions used in this study.**

| Solution name | Solution ingredients |
|---|---|
| Activated monomer solution | Sodium acrylate 19% wt/wt, acrylamide 10% v/v, BIS 0.1% v/v, in PBS, 1% w/v TEMED, 1% v/v APS |
| Denaturation buffer | 200 mM SDS, 200 mM NaCl, 50 mM Tris, pH 9, in water |
| Anchoring solution | 1.4 % v/v formaldehyde, 2 % v/v acrylamide in PBS |
| Fixation solution | 4% w/v paraformaldehyde in PBS |
| Freezing solution | 50% v/v glycerol in Milli-Q water |
| Blocking solution | 3% w/v bovine serum albumin in PBS |
| Wash buffer | 0.5% v/v TWEEN-20 in PBS |

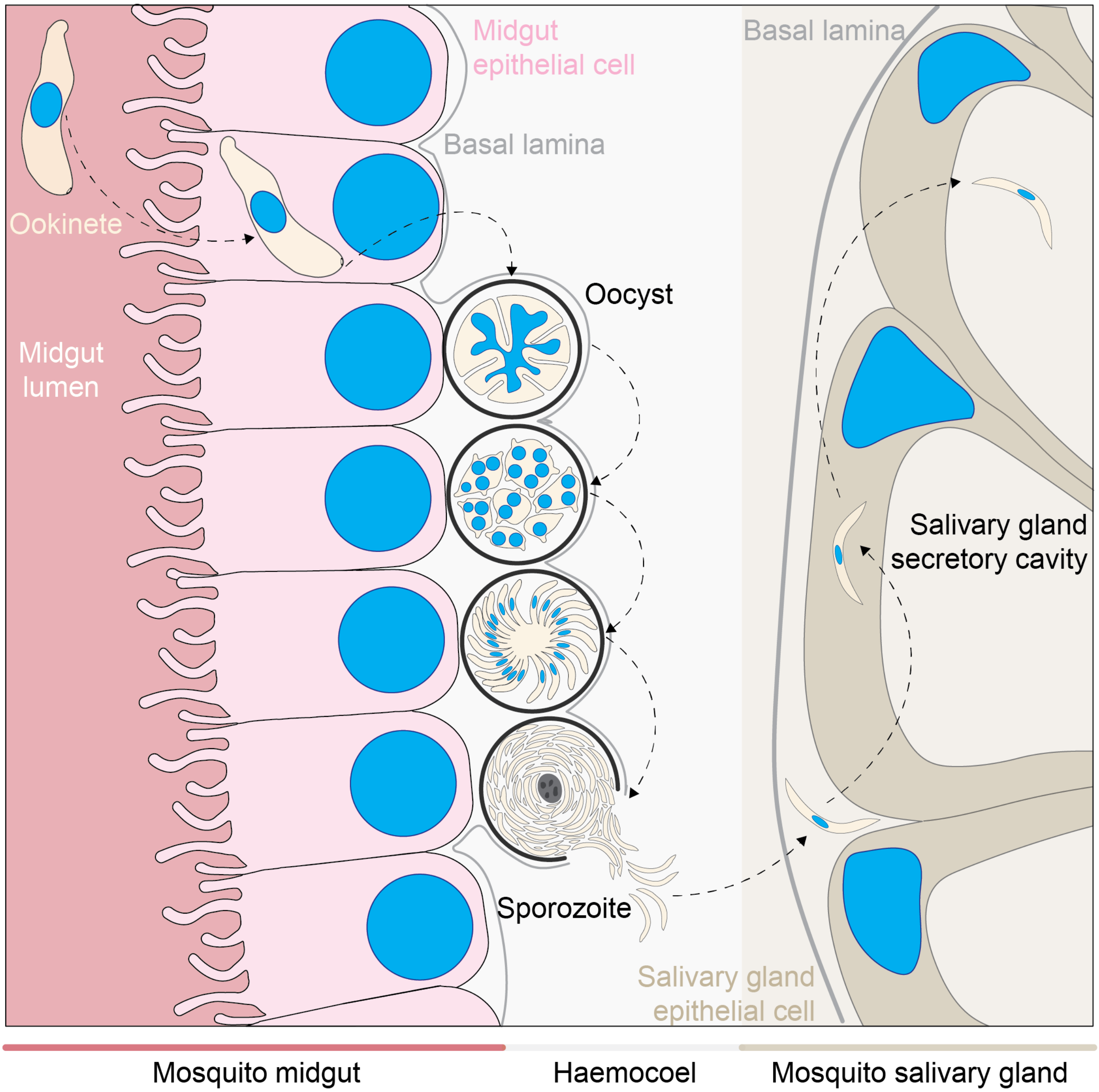


**Supplementary Figure 1: Oocyst development and sporozoite salivary gland invasion.**

Ookinetes in the midgut lumen form an oocyst on the basal side of the mosquito midgut epithelium. Over a period of several days, the oocysts undergo repeated rounds of mitosis before forming a sporoblast and beginning the process of sporozoite formation. Once sporozoites are fully mature, they will egress from the oocyst into the haemocoel of the mosquito. Haemocoel sporozoites will migrate to the salivary gland, where they first cross the basal lamina before invading the salivary gland epithelial cell and subsequently secretory cavity. Sporozoites will reside within the secretory cavity until transmitted to a host during the mosquito's next blood meal. Oocyst images adapted from[6], salivary gland invasion images adapted from[7].

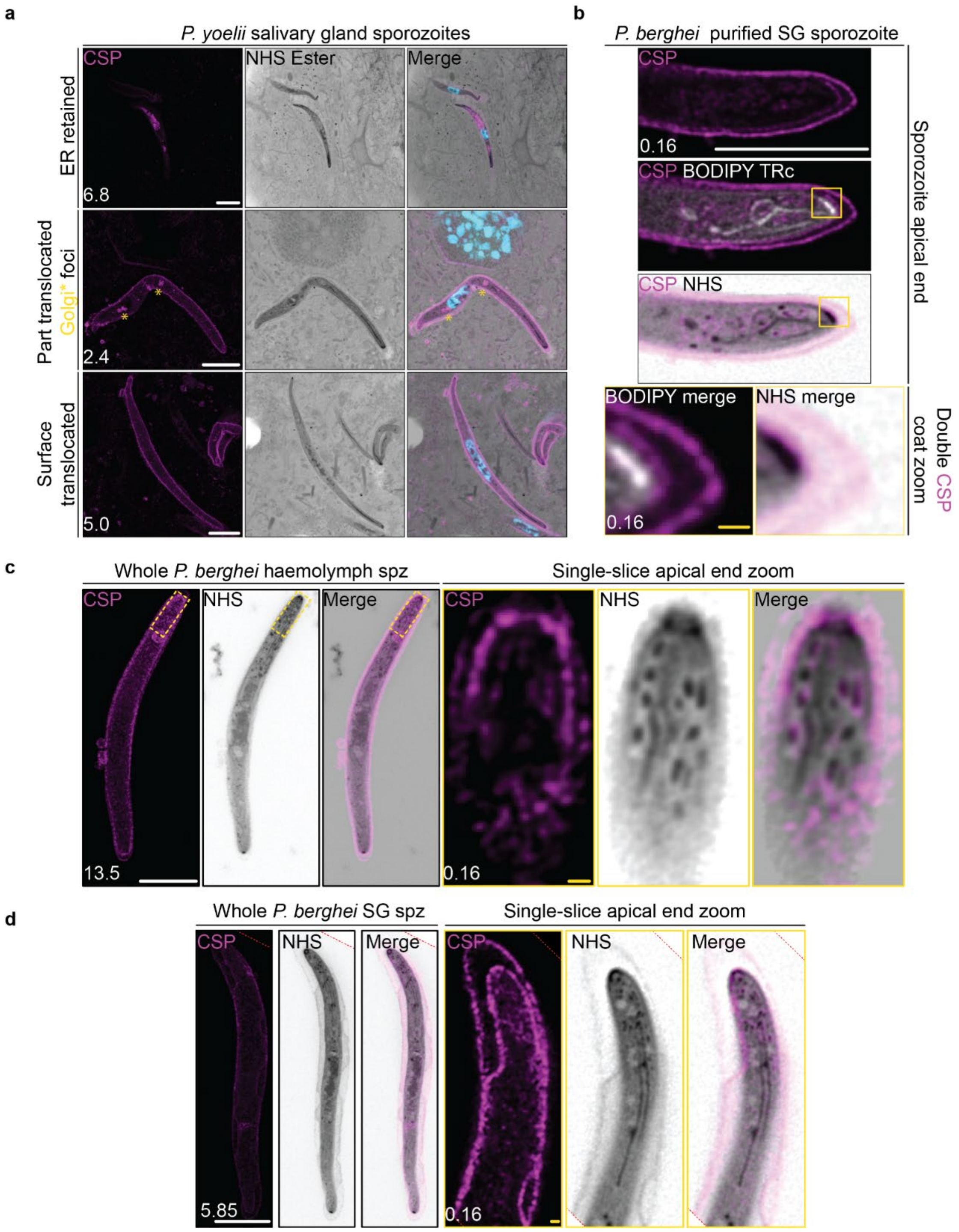


**Supplementary Figure 2: Visualisation of circumsporozoite protein in expanded sporozoites.** (**a**) *Plasmodium yoelii* infected mosquito salivary glands were prepared by MoTissU-ExM and stained with NHS Ester (protein density, greyscale), SYTOX Deep Red (DNA, cyan), and anti-circumsporozoite protein antibodies (CSP, sporozoite surface, magenta). In different sporozoites, CSP was observed at different stages of its surface translocation. Either retained predominantly in the ER, partially surface translocated with Golgi foci (yellow asterisks), or fully surface translocated. (**b**) Purified *P. berghei* SG sporozoites were prepared by U-ExM and stained as described in (**a**) but with the addition of BODIPY TR ceramide (lipids, white). Two clear layers of CSP can be observed. (**c**) *P. berghei* haemolymph sporozoites were isolated and prepared by U-ExM, stained with NHS Ester (greyscale) and anti-CSP antibodies (magenta). Protein-dense granules can be observed in the sporozoite cytosol, with some corresponding to CSP fluorescence. (**d**) As in (**c**) except for isolated *P. berghei* SG sporozoites. Scale bars: white = 10 μm, yellow = 500 nm. Number in bottom left corner indicates image Z-depth in μm.

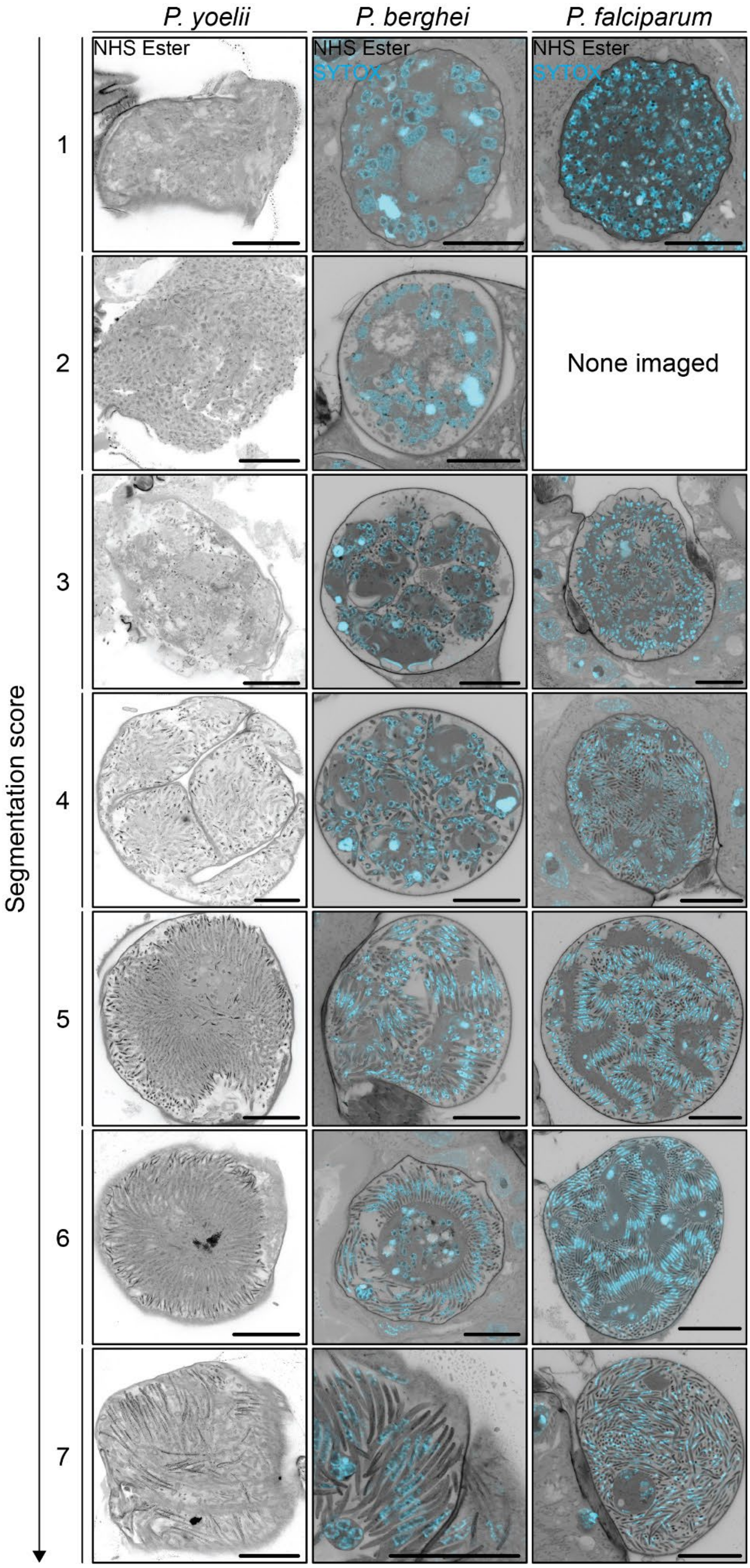


**Supplementary Figure 3: Examples of segmentation scores across *Plasmodium* spp.** Midguts infected with either *P. yoelii*, *P. berghei*, or *P. falciparum* were prepared by MoTissU-ExM and stained with NHS ester (protein density, greyscale) and SYTOX (DNA, cyan). Each image represents a single z-slice from an individual oocyst. Scale bar = 50 µm. Over the course of this study, we did not observe any *P. falciparum* oocysts with a segmentation score of 2.

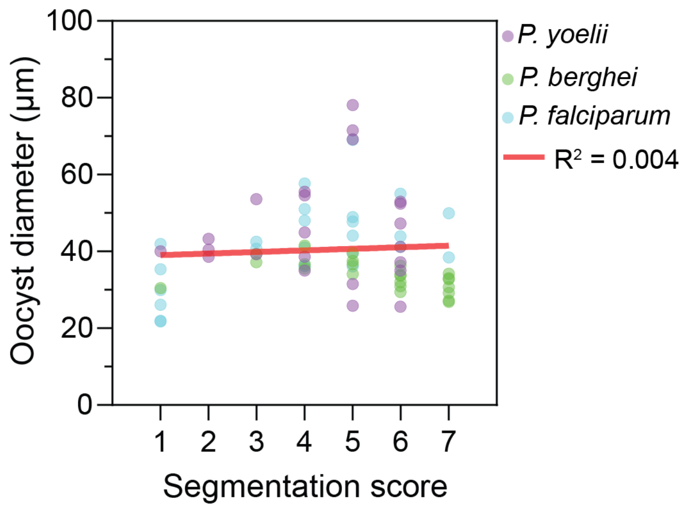


**Supplementary Figure 4: Correlation of oocyst diameter and segmentation score.** Diameter of all whole oocysts across *P. yoelii*, *P. berghei*, and *P. falciparum* imaged in this study was measured. Trendline represents simple linear regression. 71 total oocysts, (27 *Py*, 23 *Pb*, 25 *Pf*).

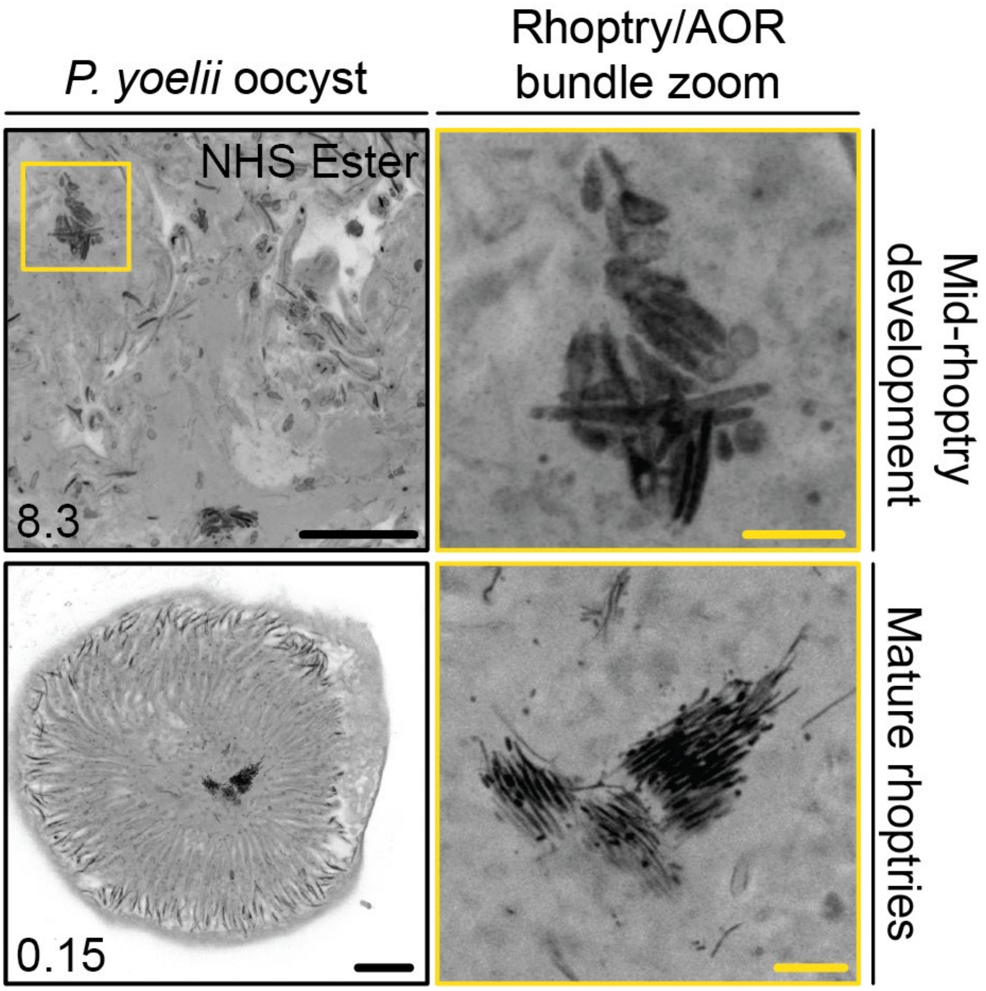


**Supplementary Figure 5: Visualisation of rhoptries and AORs outside of forming sporozoites.** Visualisation of the large bundle of rhoptries/AORs that was often found accumulating in the sporoblast, not incorporated into any of the forming sporozoites. Note that the developmental stage of the rhoptries mimic those inside the forming sporozoites. Scale bars: Black = 20 µm, yellow = 5 µm. Number in bottom left corner = image z-depth in µm.

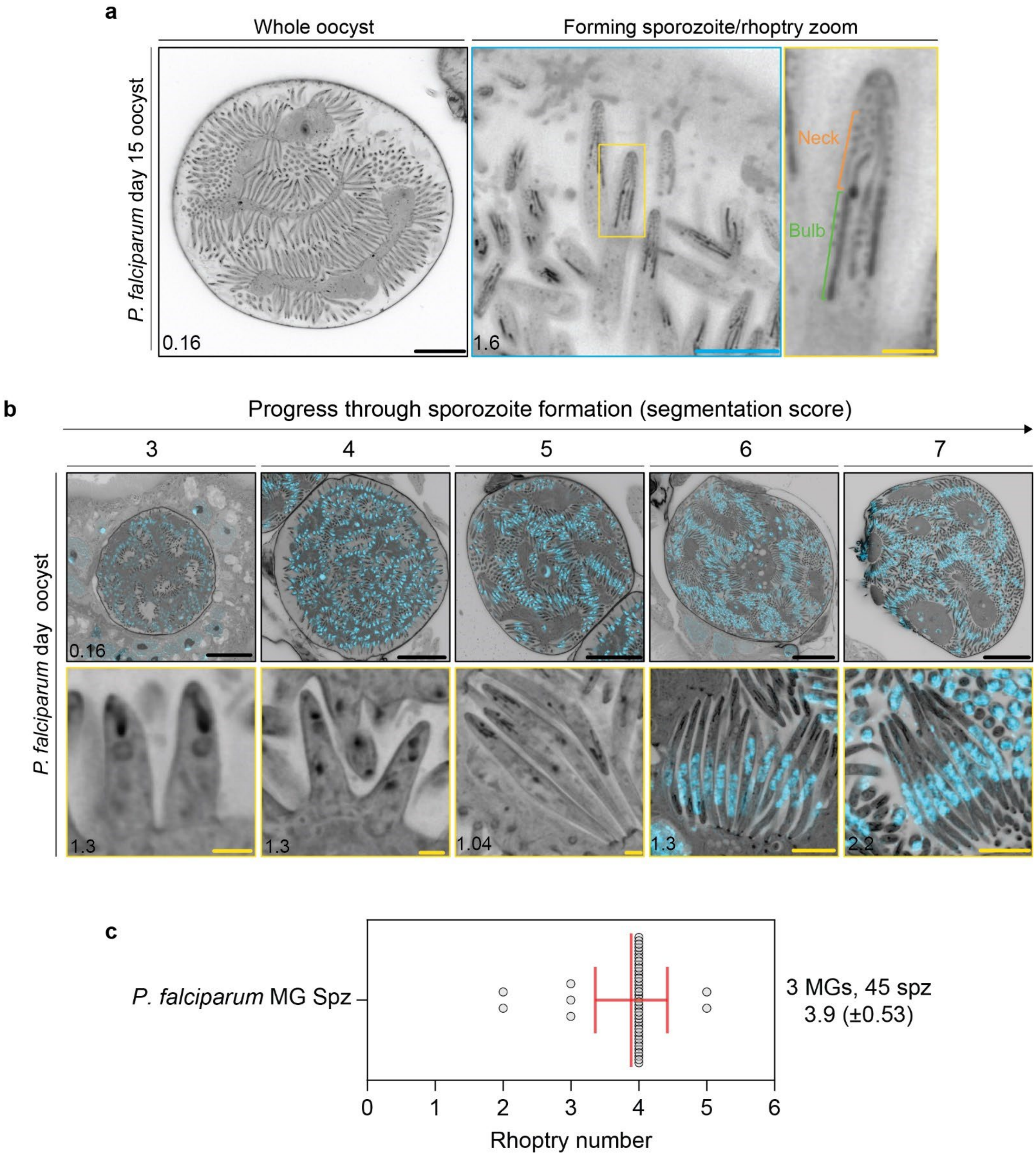


**Supplementary Figure 6: *Plasmodium falciparum* sporozoites have discernible neck and bulb regions.** Mosquito midguts at 16 days post infection with *P. falciparum* were harvested and prepared by MoTissU-ExM, stained with NHS Ester (protein density, greyscale) and SYTOX (DNA, cyan). (**a**) In mid segmentation sporozoites, rhoptries contain discernible neck and bulb regions. (**b**) Visualisation of rhoptry biogenesis during *P. falciparum* sporozoite formation. Scale bars: black = 50 µm, blue = 10 µm, yellow = 2 µm. (**c**) Quantification of rhoptry number in mature oocyst sporozoites (segmentation score 6/7). Error bars = SD.

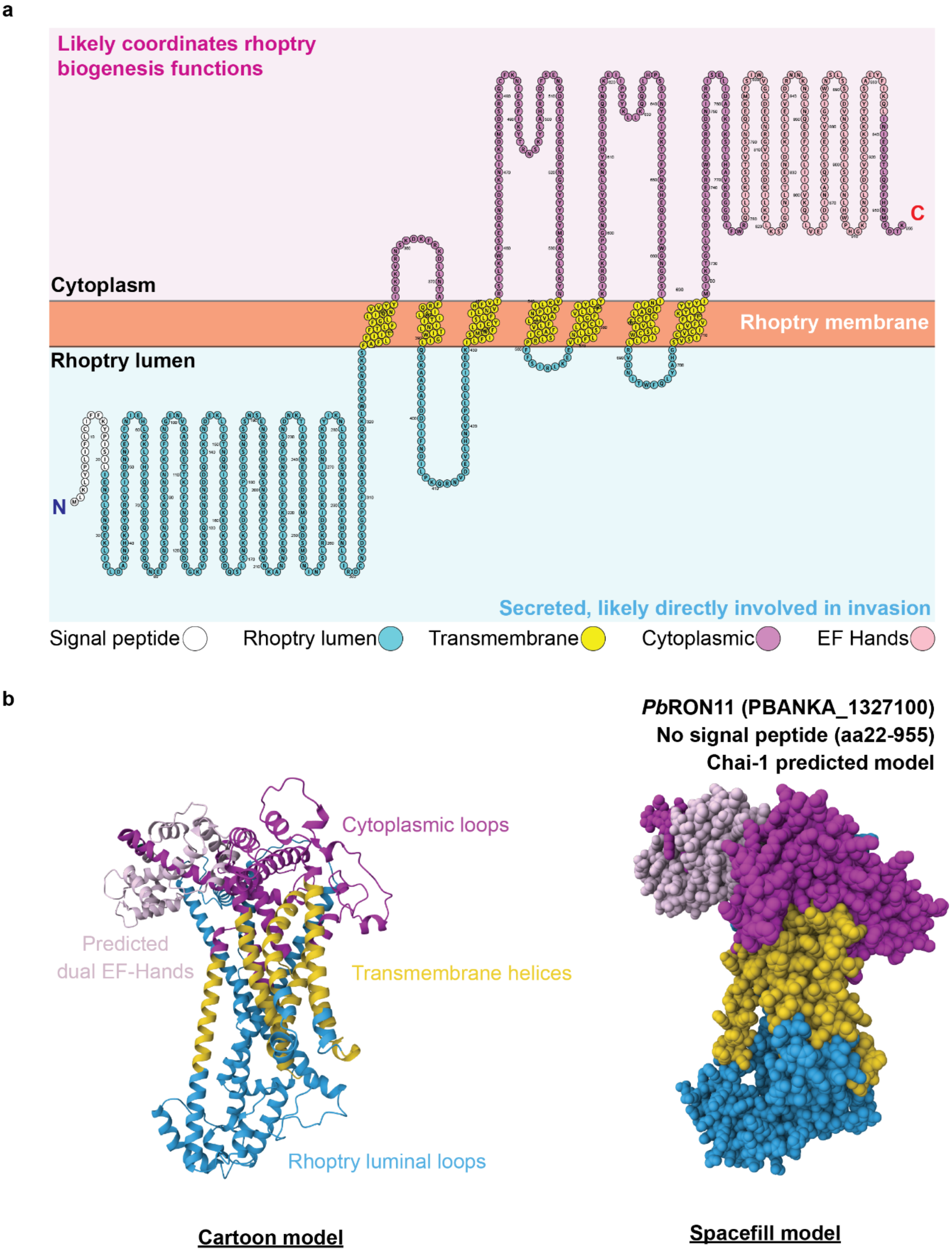


**Supplementary Figure 7: Predicted *Pb*RON11 membrane topology and protein structure.** (**a**) *Plasmodium berghei* RON11 (PBANKA_1327100) is comprised of 955 amino acids and is predicted to contain a signal peptide and 7 transmembrane regions (DeepTMHMM)[8]. This membrane topology places large regions of RON11 in both the rhoptry lumen (blue) and exposed to the cytosol (pink). Based on results from this study, and previous studies[9,10] it is likely that these regions coordinate the different functions of RON11. (**b**) The structure of *P. berghei* RON11 lacking a signal peptide (aa22-955) was predicted using Chai-1[11] and is shown as both a cartoon ribbon diagram (left) and spacefill model (right). Regions of the structure are colour coded based on their predicted membrane topology. Additionally, the location of the putative dual EF-hand domain is indicated in lilac.

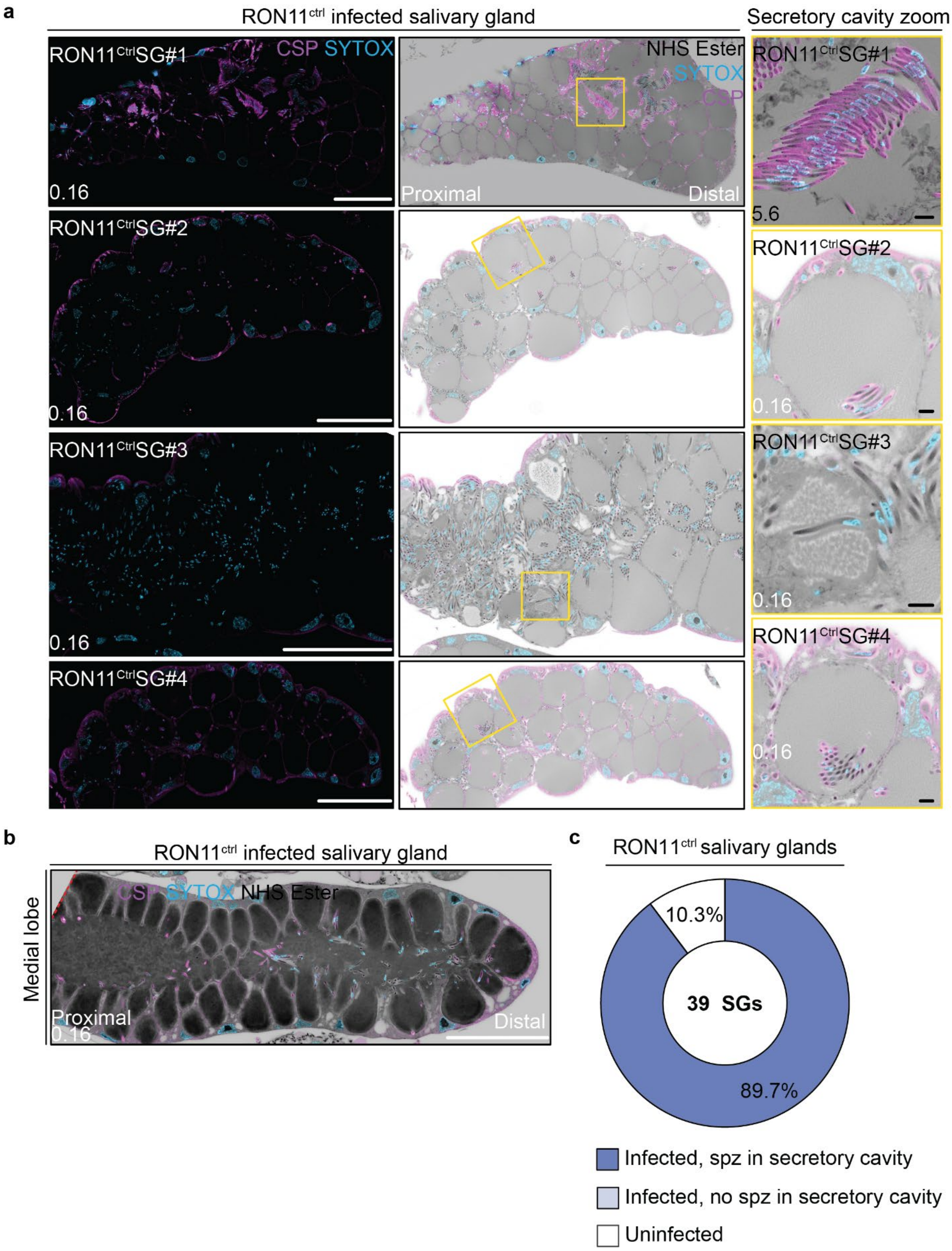


**Supplementary Figure 8: *Pb*RON11$^{ctrl}$ infected salivary glands.** Salivary glands of mosquitoes were infected with RON11$^{Ctrl}$ parasites and prepared for MoTissU-ExM, stained with the protein density dye NHS Ester (greyscale), SYTOX Deep Red (DNA, cyan) and anti-CSP antibodies (sporozoite surface, magenta). (**a**) Additional examples of RON11$^{Ctrl}$ parasites in the secretory cavity to supplement Figure 7a. (**b**) Comparison of RON11$^{Ctrl}$ infected medial salivary gland lobe. Medial lobe image was rotated for presentation purposes. Red dashed lined represents original image border. (**c**) Quantification of infection and secretory cavity invasion status of SGs for mosquitoes infected with RON11$^{Ctrl}$ parasites. Total number of SGs screened shown in centre of pie chart. Scale bars: white = 200 µm, black = 10 µm.

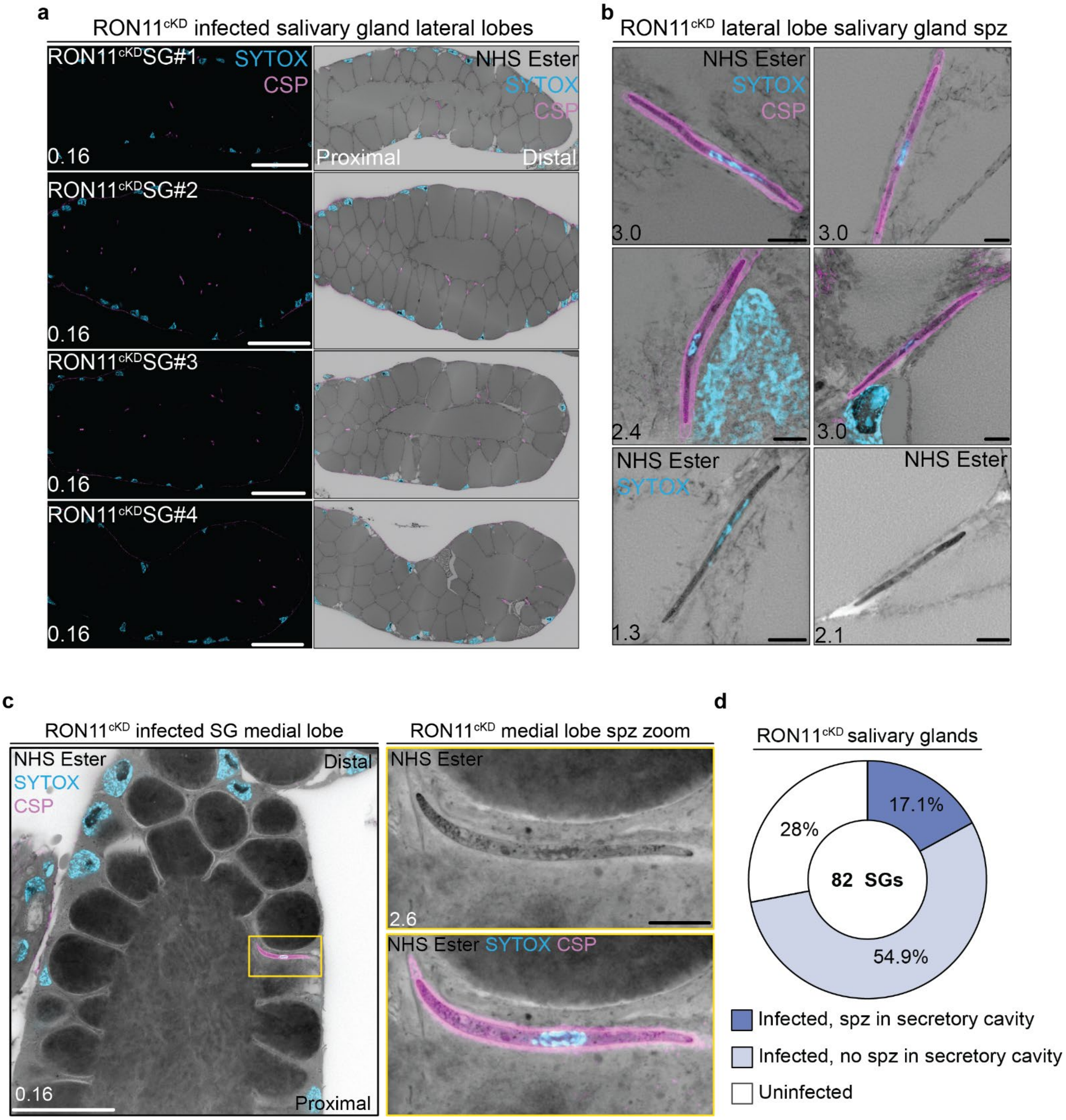


**Supplementary Figure 9: *Pb*RON11$^{cKD}$ infected salivary glands.** Salivary glands of mosquitoes were infected with RON11$^{cKD}$ parasites and prepared for MoTissU-ExM, stained with the protein density dye NHS Ester (greyscale), SYTOX Deep Red (DNA, cyan) and anti-CSP antibodies (sporozoite surface, magenta). (**a**) Further examples of RON11$^{cKD}$ infected salivary glands and (**b**) RON11$^{cKD}$ in the space between salivary gland epithelial cells (not intracellular or in secretory cavity) in addition to Figure 7 **b** and **c.** (**c**) RON11$^{cKD}$ present in the intercellular space of a medial lobe. (**d**) Quantification of infection and secretory cavity invasion status of SGs for mosquitoes infected with RON11$^{cKD}$ parasites. Total number of SGs screened shown in centre of pie chart Scale bars: white = 200 µm, black = 10 µm

**Supplementary Video 1-8: *Pb*RON11$^{cKD}$ parasites in the salivary gland intercellular space.** Salivary glands of mosquitoes were infected with RON11$^{cKD}$ parasites and prepared for MoTissU-ExM, stained with the protein density dye NHS Ester (greyscale), SYTOX Deep Red (DNA, cyan) and anti-CSP antibodies (sporozoite surface, magenta). Each video is a slice-by-slice view of a SG where RON11$^{cKD}$ sporozoites can be observed present inside the SG, but not inside the SG epithelial cell or secretory cavity.
